# The Endocytic Recycling Compartment Serves as a Viral Factory for Hepatitis E Virus

**DOI:** 10.1101/2021.10.14.464345

**Authors:** Cyrine Bentaleb, Kévin Hervouet, Claire Montpellier, Charline Camuzet, Martin Ferrié, Julien Burlaud-Gaillard, Stéphane Bressanelli, Karoline Metzger, Elisabeth Werkmeister, Maliki Ankavay, Nancy Leon Janampa, Julien Marlet, Julien Roux, Clarence Deffaud, Anne Goffard, Yves Rouillé, Jean Dubuisson, Philippe Roingeard, Cécile-Marie Aliouat-Denis, Laurence Cocquerel

## Abstract

**Background & Aims:** Although Hepatitis E virus (HEV) is the major leading cause of enterically transmitted viral hepatitis worldwide, many gaps remain in the understanding of the HEV lifecycle. Notably, viral factories induced by HEV have not been documented yet and it is currently unknown whether HEV infection leads to cellular membrane modelling as many positive-strand RNA viruses. HEV genome encodes three proteins, the ORF1 replicase, the ORF2 capsid protein and the ORF3 protein involved in virion egress. Previously, we demonstrated that HEV produces different ORF2 isoforms including the virion-associated ORF2i form. Here, we aimed to probe infectious particles and viral factories in HEV-producing cells, using antibodies directed against the different ORF2 isoforms.

**Methods:** We generated monoclonal antibodies that specifically recognize the particle-associated ORF2i form, and antibodies that recognize the different ORF2 isoforms. We used them in confocal and electron microscopy approaches to probe viral factories in HEV-producing cells. We performed an extensive colocalization study of viral proteins with subcellular markers. We analyzed the impact of silencing Rab11, a central player of the endocytic recycling compartment (ERC).

**Results:** One of the antibodies, named P1H1 and targeting the N-terminus of ORF2i, recognized delipidated HEV particles. Confocal and ultrastructural microscopy analyses of HEV-producing cells revealed an unprecedented HEV-induced membrane network containing tubular and vesicular structures. These subcellular structures were enriched in ORF2 and ORF3 proteins, and were dependent on the ORF3 expression and ORF2i capsid protein assembly. Colocalization and silencing analyses revealed that these structures are derived from the ERC.

**Conclusions:** Our study reveals that HEV hijacks the ERC and forms a membrane network of vesicular and tubular structures that might be the hallmark of HEV infection.

**Lay summary:** Hepatitis E virus (HEV) is the leading cause of acute hepatitis worldwide but many steps of its lifecycle are still elusive. Thanks to the development of new antibodies that recognize the different forms of the HEV capsid protein, we were able to visualize vesicular and tubular structures that were established by the virus in the host cell. In addition, extensive efforts to identify these structures led us to conclude that HEV hijacks the endocytic recycling compartment of the cell to form this network of vesicles and tubules, which might be the hallmark of HEV infection.

## Introduction

Hepatitis E virus (HEV) is the most common cause of acute viral hepatitis worldwide. Four main distinct genotypes (gt), belonging to a single serotype, infect humans. HEV gt1 and gt2 are restricted to humans and are responsible for waterborne outbreaks in developing countries with low sanitary conditions. HEV gt3 and gt4 are zoonotic and cause sporadic zoonotic foodborne hepatitis in industrialized countries [1–3]. Although HEV causes a mostly asymptomatic self-limited disease, gt1-infection can lead to fulminant liver failure, particularly in pregnant women, and gt3-infection can lead to chronic disease in immunosuppressed patients. There is no specific treatment nor universal vaccine against HEV [4].

HEV is found as a non-enveloped virus in bile and feces or as a quasi-enveloped virus (eHEV) in blood and cell culture. Its RNA genome encodes three proteins, the ORF1 replicase, the ORF2 capsid protein and the ORF3 protein involved in virion egress [5]. Previously, we demonstrated that, during the HEV lifecycle, HEV produces at least 3 forms of the ORF2 capsid protein: infectious ORF2 (ORF2i), glycosylated ORF2 (ORF2g), and cleaved ORF2 (ORF2c) [6]. The ORF2i protein is the structural component of infectious particles that is likely derived from the assembly of the intracellular ORF2i form. In contrast, ORF2g and ORF2c proteins (herein referred to as ORF2g/c) are not associated with infectious material but secreted in large amounts (*i.e.,* about 1000x more than ORF2i [7]) and are the most abundant antigens detected in patient sera [6]. In addition, these proteins likely act as humoral decoys that inhibit antibody-mediated neutralization [7]. Recently, we demonstrated that a 5 amino acid Arginine-Rich Motif (ARM) located in the ORF2 N-terminal region is a unique central regulator of ORF2 addressing that finely controls the HEV lifecycle [8]. Indeed, the ARM controls ORF2 nuclear translocation, promoting regulation of host antiviral responses. This motif also regulates the dual topology and functionality of ORF2 signal peptide, leading to the production of either cytosolic infectious ORF2i or reticular non- infectious glycosylated ORF2 forms. Furthermore, the ARM likely serves as a cleavage site of the glycosylated ORF2 protein. Finally, it promotes ORF2 membrane association that is likely essential for particle assembly [8].

Recent breakthroughs have been achieved in developing cell culture models for HEV [9]. However, many gaps remain in the knowledge of the HEV lifecycle such as the intracellular location of HEV replication and particle assembly as well as the underlying mechanisms of these processes [10]. It is known that the majority of positive sense single-stranded RNA viruses induce host cell membrane rearrangements to facilitate their viral genome replication, viral particle assembly and to protect from the innate immune response. These membrane rearrangements have been well-characterized by confocal and electron microscopy approaches leading to the identification of a broad spectrum of complexity between the host membrane remodeling and viral and cellular actors involved in these arrangements [11]. However, due to a scarcity of robust cell culture models, membrane remodeling and viral factories induced by HEV have not been documented yet. Here, we generated monoclonal antibodies directed against the ORF2 capsid protein and used them to probe infectious particles and viral factories in HEV-producing cells.

## Materials and methods

### Antibodies

Mice immunization and generation of P1H1, P2H1, P2H2 and P3H2 monoclonal antibodies were performed as described in [12] and in **supplementary information**. Other primary antibodies used in this study are listed in **Supplementary Table 1**. Peroxidase- and fluorochrome-conjugated secondary antibodies were from Jackson ImmunoResearch. Gold-conjugated secondary antibodies were from Aurion (Wageningen, Netherlands).

### Cells

PLC3 cells are a subclone of PLC/PRF/5 hepatoma cells [6]. PLC3 and Huh-7.5 [13] cells were authenticated by STR profiling and Multiplex Cell Authentication (Multiplexion) respectively, and cultured as previously described [14].

### Plasmids and transfection

The plasmid pBlueScript SK(+) carrying the DNA of the full length genome of gt3 Kernow C-1 p6 strain, (GenBank accession number JQ679013, kindly provided by S.U Emerson) was used [15]. The ORF3-null mutant (HEV-p6-ΔORF3) and the 5R/5A mutant (HEV-p6-5R/5A) of HEV-p6 were generated as described in [16] and [8], respectively.

The plasmid pBlueScript SK(+) carrying the DNA of the p6-V1-Puro replicon was generated from the pBlueScript SK(+)/p6-V1 plasmid [17] in which the ORF3/ORF2 region was replaced by a fragment encoding the resistance gene to puromycin as well as the last 321 amino acid residues of ORF2.

Capped genomic HEV RNAs were prepared with the mMESSAGE mMACHINE kit (Ambion) and delivered to PLC3 or Huh-7.5 cells by electroporation using a Gene Pulser Xcell^TM^ apparatus (Bio-Rad) [6]. PLC3/HEV and Huh-7.5/HEV cells were electroporated with p6 RNA (20μg / 4.10^6^ cells) whereas PLC3 mock and Huh-7.5 mock were electroporated in the absence of RNA.

### PLC3-replicon

Cells stably harboring a p6 subgenomic replicon were generated by electroporating PLC3 cells with p6-V1-Puro replicon capped RNAs. Cells were next cultured with 2.5 µg/ml of puromycin (Euromedex) for the selection of cells transfected with the ORF1-V5-puro replicon.

### Indirect immunofluorescence

Immunostainings were performed as described in **supplementary information.** For high-resolution confocal analyses, images were acquired with an Airyscan module. Z-stacks acquisition were performed with a 0.15 μm z-interval. The ORF2 confocal parameters were determined to optimize the dynamic range and avoid intensity saturation, and the same settings were applied for each sample. Images in the 3 channels were recorded using Zen image collection software (Carl Zeiss Microscopy) and processed for high resolution reconstruction. 3D volumetric surface constructs were obtained using Imaris software (version 9.5.1; Oxford instruments, Belfast), by applying an intensity threshold for each channel. The images were then processed using Fiji software.

### Manders’ overlap coefficient (MOC) determination

Colocalization studies were performed by calculating the MOC using the JACoP plugin of Fiji software. For each calculation, at least 30 cells were analyzed to obtain a MOC mean. A MOC of 1 indicates perfect overlap and 0 no overlap.

### Western blotting analysis

Western blotting analyses were performed as described in **supplementary information**. Target proteins were detected with specific antibodies (**Supplementary Table 1**) and corresponding peroxidase-conjugated secondary antibodies.

### Immunoprecipitations (IP)

Antibodies were bound to Tosylactivated M-280 Dynabeads^TM^ (Thermofisher) overnight at 37°C following the manufacturer’s recommendations. Beads were washed and then incubated for 1h at room temperature with supernatants (heat-inactivated or Triton-X100-treated), lysates or Triton-X100- treated patient sera. Beads were washed and then heated at 80°C for 20 min in Laemmli buffer. ORF2 proteins were detected by WB using the 1E6 antibody.

### qIP and RT-qPCR

For qIP experiments, samples were first immunoprecipitated as described above. Next, RNAs were extracted and quantified by RT-qPCR as described in **supplementary information**.

### Immunoelectron microscopy

Electroporated cells were fixed by incubation for 2 h with 4% paraformaldehyde in phosphate buffer (pH 7.6), washed twice with PBS (pH 7.6), and centrifuged at 300 × g for 10 min. Cell pellets were embedded in 12% gelatin and infused with 2.3 M sucrose overnight at 4°C. Ultra-thin cryosections (90 nm) were cut at −110°C on a LEICA FC7 cryo-ultramicrotome. The sections were retrieved in a mixture of 2% methylcellulose and 2.3 M sucrose (1:1) and collected onto formvar/carbon-coated nickel grids. The gelatin was removed by incubation at 37°C, and the sections were incubated with primary antibodies (**Supplementary Table 1**). The grids were washed with PBS and then incubated with secondary antibodies conjugated to gold particles of 6 nm or 10 nm in diameter. The grids were washed with PBS, post-fixed in 1% glutaraldehyde and rinsed with distilled water. Contrast staining was achieved by incubating the grids with a 2% uranyl acetate/2% methylcellulose mixture (1:10). The sections were then examined under a transmission electron microscope operating at 100 keV (JEOL 1011). Electron micrographs were recorded with a digital camera driven by Digital Micrograph software (GMS 3, Gatan).

### Silencing experiments

At 3 days p.e., PLC3/HEV cells were transfected with small interfering RNA (siRNA) pools (Horizon) targeting Rab11a (ON-TARGETplus human Rab11a, gene 8766, siRNA SMARTpool) and Rab11b (ON-TARGETplus human Rab11b, gene 9230, siRNA SMARTpool) (siRab11) or with a nontargeting siRNA control (siCTL) by using RNAiMax reagent (Invitrogen) according to the manufacturer’s instructions. The knockdown effects were determined at 72h post-transfection by immunofluorescence, western-blotting, RT-qPCR and virus titration.

## Results

### Generation of monoclonal antibodies that specifically recognize the ORF2i protein

The ORF2 protein sequence contains 660 amino acids (aa) (**Fig. 1A**). Previously, we demonstrated that the first aa of ORF2i, ORF2g and ORF2c proteins are L^14^, S^34^ and S^102^, respectively [6, 14]. The ORF2i protein is not glycosylated whereas ORF2g/c proteins are N-glycosylated on ^137^NLS and ^562^NTT sequons [14] (**Fig. 1A**). We capitalized on these features to design two immunogen peptides (P1 and P2) for obtaining highly specific antibodies of the ORF2i form. The P1 peptide corresponds to the N-terminus of the ORF2i protein that is not present in the ORF2g/c form sequences. The P2 peptide corresponds to 14 aa covering the ^562^NTT sequon that is not occupied by N-glycans on the ORF2i protein, in contrast to ORF2g/c proteins. We also designed a P3 peptide for obtaining antibodies recognizing the different forms of ORF2 (**Fig. 1A**). To verify the specificity between strains/genotypes, sequence alignments were carried out for each peptide (data not shown). Following mice immunization and hybridoma screening, one clone from P1 immunization (P1H1), two clones from P2 immunization (P2H1 and P2H2) and one clone from P3 immunization (P3H2) were selected and further characterized.

**Fig 1:**
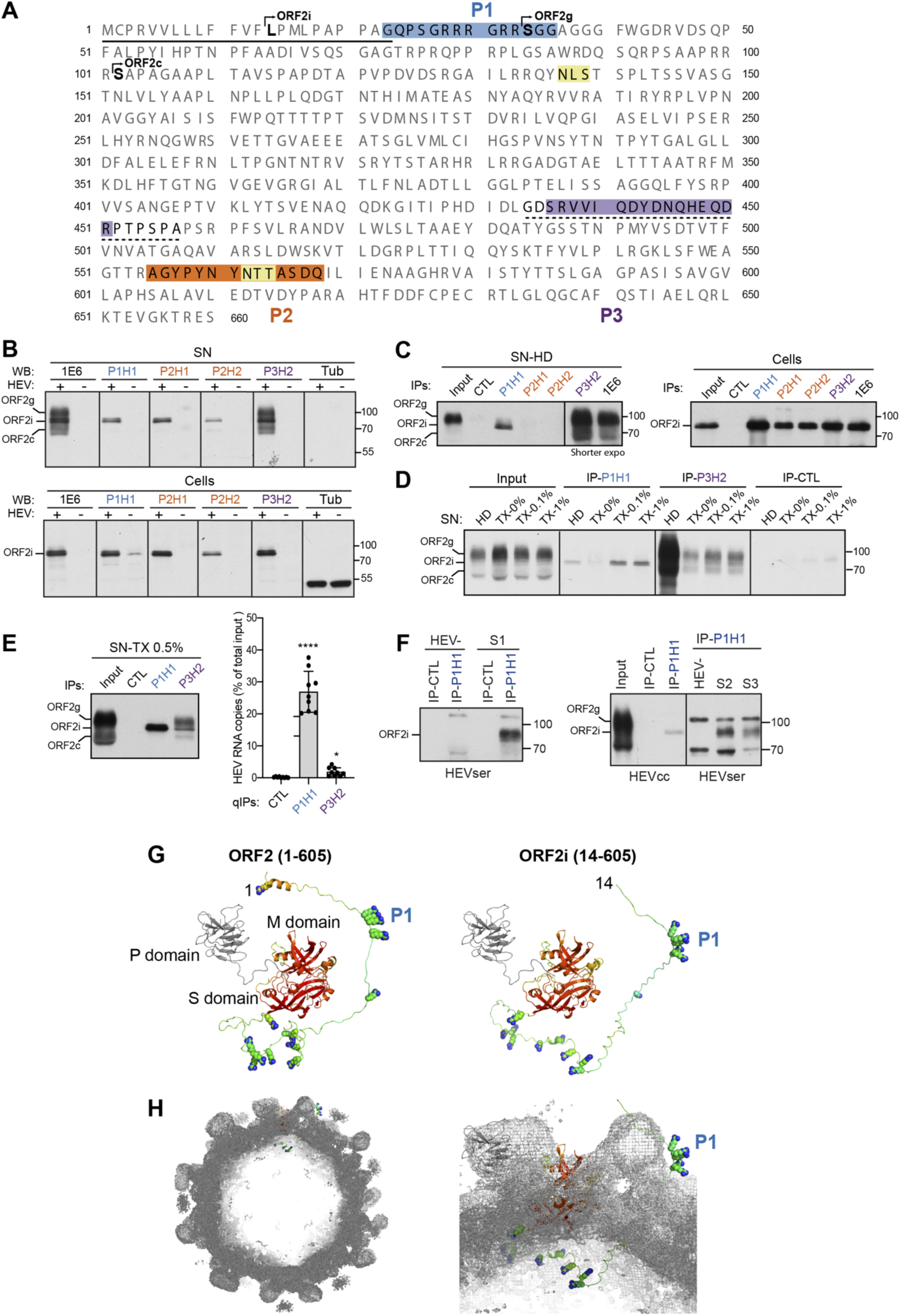
Generation of monoclonal antibodies that specifically recognize the ORF2i protein. (**A**) Sequence of ORF2 proteins. The line corresponds to the signal peptide. N-glycosylation sites are highlighted in yellow. P1, P2 and P3 peptides are highlighted in blue, orange and purple, respectively. The dashed line corresponds to the 1E6 epitope. (**B**) Detection of ORF2 proteins in supernatants (SN) and lysates (cells) of PLC3/HEV cells (+) and non-infected PLC3 cells (-) by WB. (**C**) Immunoprecipitation of ORF2 proteins in heat-denatured (HD, 20min at 80°C) SN and lysates of PLC3/HEV cells. An anti-HCV E2 envelope protein antibody (H52) was used as a negative control (CTL). ORF2 proteins were detected by WB with the 1E6 antibody (WB 1E6). (**D**) IP of SN treated for 30min with Triton X-100 (TX-0.1%, TX-1%), or left untreated (TX-0%). Input used for IP are shown on the left. ORF2 proteins were detected by WB 1E6. (**E**) SN of PLC3/HEV cells was treated with TX-0.5% and immunoprecipitated with the P1H1, P3H2 or isotype control antibodies. Half of the IP was analyzed by WB 1E6 (left panel) and the other half was processed for RNA extraction and HEV RNA quantification (right panel). Results are expressed as percentage of immunocaptured HEV RNA copies compared to the total input. (**F**) Sera of HEV-infected (HEVser, S1- S3) or non-infected (HEV-) patients were treated with TX-0.5% and immunoprecipitated with P1H1 or isotype control antibodies. IPs on SN of PLC3/HEV cells (HEVcc) were used as controls. ORF2 proteins were detected by WB with the 1E6 antibody (WB 1E6). (B-F) Molecular mass markers are indicated on the right (kDa). For clarity and conciseness concerns, blots were cropped. **(G)** Models of ORF2 (left) and ORF2i (right) including their N-termini (residues 1-128 and 14-128, respectively. The first residue is labeled. The 606-660 residues located downstream of the P domain are not included). Models are displayed in ribbons representation and, for S (shell) and M (middle) domains, colored by expected accuracy from red (most accurate) to blue (least accurate or no defined structure). P (protruding) domain is colored in gray. Arginine side chains upstream residue 128 are displayed as spheres. The P1 epitope is labeled. (**H**) The same ORF2i model in the same representation and orientation as in (G, right) has been fitted at the ’A’ molecule position in the 12-Å cryo- EM map of a virion-sized recombinant ORF2 icosahedral T=3 particle (EMDB 5173). Left, cutaway overall view. Right, zoom on the model.

Specificity of generated antibodies was first analyzed in western blotting (WB) experiments with supernatants (SN) and lysates of PLC3 cells electroporated with the gt3 p6 strain (PLC3/HEV cells) and non-infected PLC3 cells (**Fig. 1B**). SN of PLC3/HEV cells contains quasi-enveloped infectious HEV particles (ORF2i) but also large amounts of ORF2g/c proteins, whereas cells express the intracellular ORF2i form that assembles to form intracellular particles [6, 14]. The 1E6 monoclonal antibody, which recognizes the three forms of ORF2 proteins [6] was used as a control. The P1H1, P2H1 and P2H2 antibodies showed a highly specific recognition of the ORF2i protein without cross-reacting with the ORF2g/c proteins, as compared to the P3H2 and 1E6 antibodies that recognized the three forms (**Fig. 1B**).

Antibodies were next used in immunoprecipitation (IP) assays against heat-denatured (HD) SN and lysates of PLC3/HEV cells. In lysates, all antibodies immunoprecipitated the ORF2i protein (**Fig. 1C**, right panel). In contrast, in SN, no ORF2 proteins were detected in IP-P2H1 and IP-P2H2 whereas the IP-P1H1 showed a highly specific recognition of the ORF2i protein, without cross-reaction with the ORF2g/c proteins (**Fig. 1C**, left panel), as compared to IP-P3H2 and IP-1E6 that displayed the three forms of ORF2. Thus, P1H1 specifically immunoprecipitates heat-denatured HEV particles.

HEV particles produced in cell culture are wrapped with lipids that mask the ORF2 epitopes. In order to analyze the ability of antibodies to immunoprecipitate non- denatured particles, SN of PLC3/HEV cells was then treated with Triton X-100 (TX), or left untreated (TX-0%) before IP with P1H1 and P3H2 (**Fig. 1D**). In untreated samples, no ORF2 proteins were detected in IP-P1H1, whereas ORF2g/c proteins were immunoprecipitated by P3H2. In treated samples, the IP-P1H1 showed a highly sensitive and specific recognition of the ORF2i protein, whereas IP-P3H2 displayed the three forms of ORF2. Thereby, P1H1 specifically immunoprecipitates detergent- treated HEV particles.

We next analyzed by RT-qPCR the ability of P1H1 and P3H2 antibodies to immunocapture HEV particles in TX-treated SN. We found that P1H1 immunocaptured 27% of total RNA input whereas P3H2 immunocaptures only 2% (**Fig. 1E**). These results indicate that P1H1 efficiently recognizes non-lipid-associated-particles and that its epitope is likely exposed on naked HEV particles.

Importantly, we analyzed the ability of P1H1 antibody to recognize particles from infected patient sera (HEVser). For this purpose, IP-P1H1 and IP-CTL were performed on TX-0.5%-treated sera from HEV-infected (S1-S3) and non-infected (HEV-) patients. As shown in **Figure 1F**, P1H1 efficiently immunoprecipitated the particle-associated ORF2i form, indicating that the P1H1 antibody captures patient HEV particles.

Finally, in order to visualize how the P1 epitope may be exposed on HEV particles, we modelled the N-terminal part of ORF2. The N-terminus (residues 1-128 for ORF2 and 14-128 for ORF2i) was clearly flexible and found in different places in different models. However, the very first residues, including P1, were often found on the same level as or even above domains M and P (**Fig. 1G**), that are exposed at the surface of the icosahedral HEV capsid. Indeed, our models with the most extended N-termini, when placed in the context of the available 12-Å cryo-EM map of the recombinant ORF2 particle [18], readily display the P1 peptide. It is noteworthy that in all models, arginine residues of the N-terminus downstream P1 were still exposed on the inside of the capsid (**Fig. 1H**).

Antibodies were also tested for their ability to neutralize non-enveloped particles but none of them displayed neutralizing activity (data not shown), indicating that, while exposed on HEV particles, the P1 epitope is not involved in HEV entry.

### Identification of HEV-induced subcellular structures

We next performed double-label immunofluorescence and confocal analyses of PLC3/HEV (**Fig. 2A**) and PLC3 mock (**Fig. S1**) cells using the anti-ORF2 antibodies and a polyclonal antibody directed against ORF3, a small protein with a viroporin activity [19] that is associated with the quasi-enveloped viral particle and supports virion egress through the exosomal pathway [20, 21]. ORF3 protein is palmitoylated at cysteine residues in its N-terminal region and is exposed to the cytosolic side of the membrane [22]. The anti-ORF2i antibodies (P1H1, P2H1 and P2H2) mainly displayed a perinuclear nugget-like ORF2 staining, whereas the P3H2 and 1E6 antibodies showed a diffuse ER-like ORF2 staining in addition to the perinuclear nugget-like ORF2 staining (**Fig. 2A**). Of note, P3H2 and 1E6 antibodies recognize the different ORF2 isoforms notably the ORF2g/c forms which are proteins going through the secretory pathway [6,8,14].

**Fig 2:**
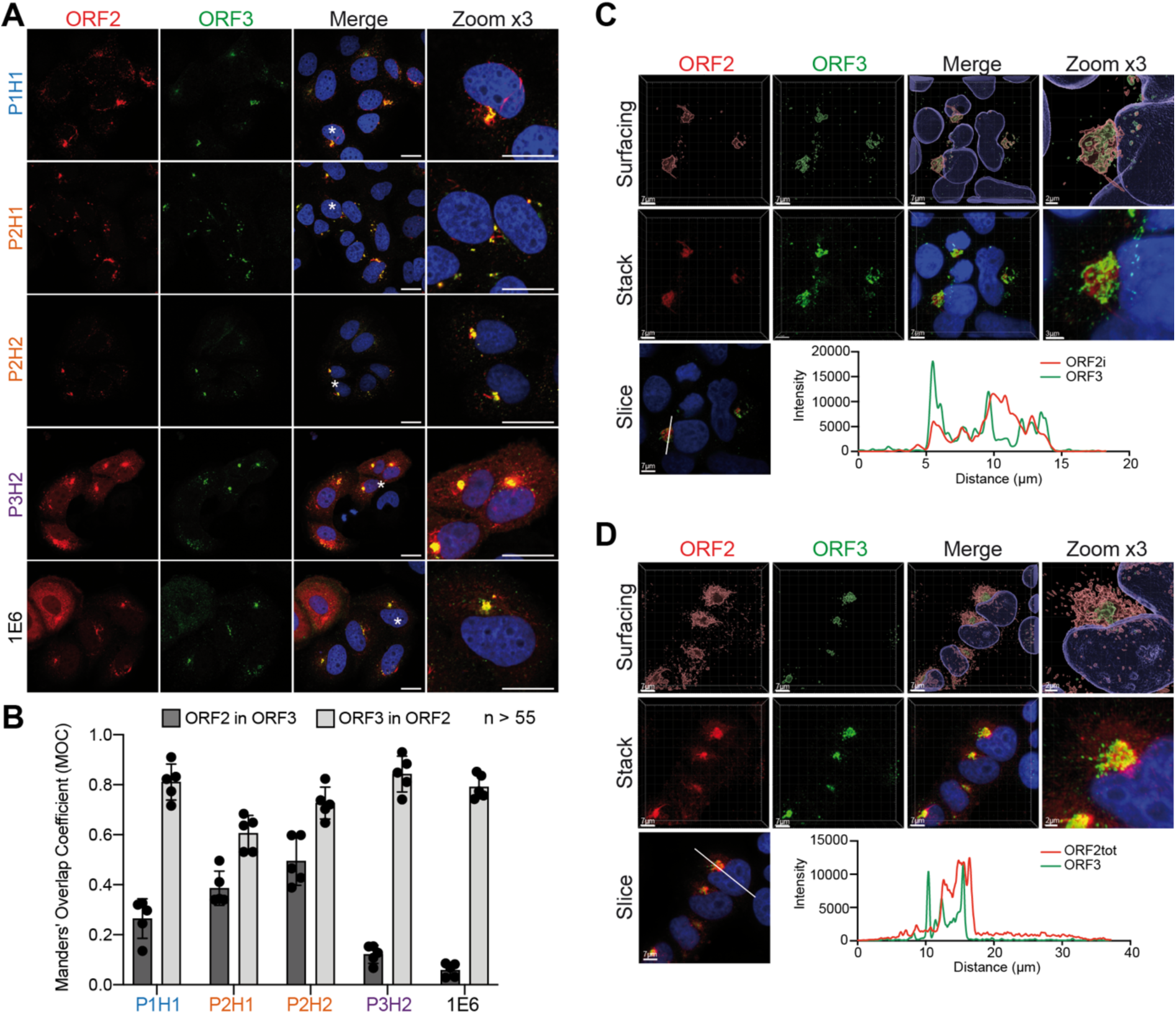
Subcellular structures recognized by anti-ORF2 antibodies. (**A**) At 6 days post-electroporation (d.p.e.), PLC3/HEV cells were fixed, permeabilized with cold methanol and TX-0.5% and double-stained with indicated anti-ORF2 and anti-ORF3 antibodies. Red = ORF2; Green = ORF3; Blue = DAPI. Staining were analyzed by confocal microscopy. Scale bar, 20μm. (**B**) Manders’ Overlap Coefficients (MOC) of the ORF2 labeling in the ORF3 labeling (ORF2 in ORF3, dark grey) and MOC of the ORF3 labeling in the ORF2 labeling (ORF3 in ORF2, light grey). (**C-D**) PLC3/HEV cells were co-stained with anti-ORF3 and P1H1 (**C**) or P3H2 (**D**) antibodies and analyzed by confocal microscopy with a high resolution Airyscan module. On the top, volume rendering of the 3D z-stacks (Surfacing) using Imaris are shown to visualize the ORF2/ORF3 substructures. In the middle, z-stacks are shown. On the bottom, line graphs show the fluorescence intensities of ORF2 and ORF3 staining measured every 50 nm across the region of interest highlighted by the white line in the micrograph shown on the bottom left of each panel. Scale bars show the indicated length.

For each antibody, we calculated the Manders’ Overlap Coefficient (MOC) of either ORF2 staining in ORF3 staining (**Fig. 2B**, ORF2 in ORF3) or ORF3 staining in ORF2 staining (**Fig. 2B**, ORF3 in ORF2). We found that the ORF3 protein staining highly overlapped with ORF2 for all antibodies, indicating that the ORF3 protein highly colocalizes with the ORF2 proteins (**Fig. 2B**, MOC ORF3 in ORF2). Moreover, the P1H1, P2H1 and P2H2 antibodies showed a higher MOC of ORF2 in ORF3 than that of P3H2 and 1E6, indicating that ORF3 protein mainly colocalizes with the ORF2i protein (**Fig. 2B**, ORF2 in ORF3). In line with this, super-resolution confocal microscopy analyses of PLC3/HEV cells stained with either P1H1 (ORF2i, **Fig. 2C**) or P3H2 (ORF2tot, **Fig. 2D**) showed a total overlap of fluorescence intensities of ORF2i with ORF3 (**Fig. 2C**) whereas a shift of fluorescence intensities was observed between total ORF2 and ORF3 proteins (**Fig. 2D**).

Together, these results indicate that anti-ORF2 antibodies, and more specifically anti- ORF2i antibodies, recognize perinuclear ORF2-enriched structures that are in close proximity to the ORF3 protein.

Of note, all antibodies recognized ORF2 proteins from HEV-gt1 (Supplementary data, **Fig. S2**).

We then performed electron microscopy (EM) experiments, using immunogold labeling with P1H1, P2H2, P3H2 and 1E6 anti-ORF2 antibodies (visualized by 6 nm gold particles) of ultrathin cryosections of PLC3/HEV (**Fig. 3A)** and PLC3 mock (**Fig. S3**) cells. On cryosections of PLC3/HEV cells, the four antibodies specifically labeled two types of subcellular structures *i.e.* tubular structures and vesicular structures (**Fig. 3A).** Tubular structures displayed a homogeneous diameter of 20-25 nm. They were highly organized, often arranged in compact parallel arrays. Vesicular structures were larger and heterogeneous in size with a diameter of 50-250 nm. Both structures were mostly found in the vicinity of the nucleus and led to a nuclear deformation (**Fig. 3A**, P3H2, asterisk). In most cells, we observed an extensive membrane network containing tubular and vesicular structures (**Fig. 3B**, P1H1, P2H2 and 1E6) and in some cells, transverse and longitudinal sections of tubular structures were observed (**Fig. 3B**, P3H2). These results suggest that the perinuclear ORF2-enriched structures previously observed in confocal microscopy likely correspond to the tubular and vesicular structures observed in EM. Importantly, HEV-induced tubular and vesicular structures were also observed in Huh-7.5 cells electroporated with HEV RNA (**Fig. S4A-C**) or infected with HEV particles (**Fig. S4D-F**). These structures were not found in cryosections of PLC3 and Huh-7.5 mock cells (**Fig. S3**). Together, these results indicate that, during its lifecycle, HEV induces the formation of perinuclear ORF2- enriched ultrastructures in the host cell.

**Fig 3:**
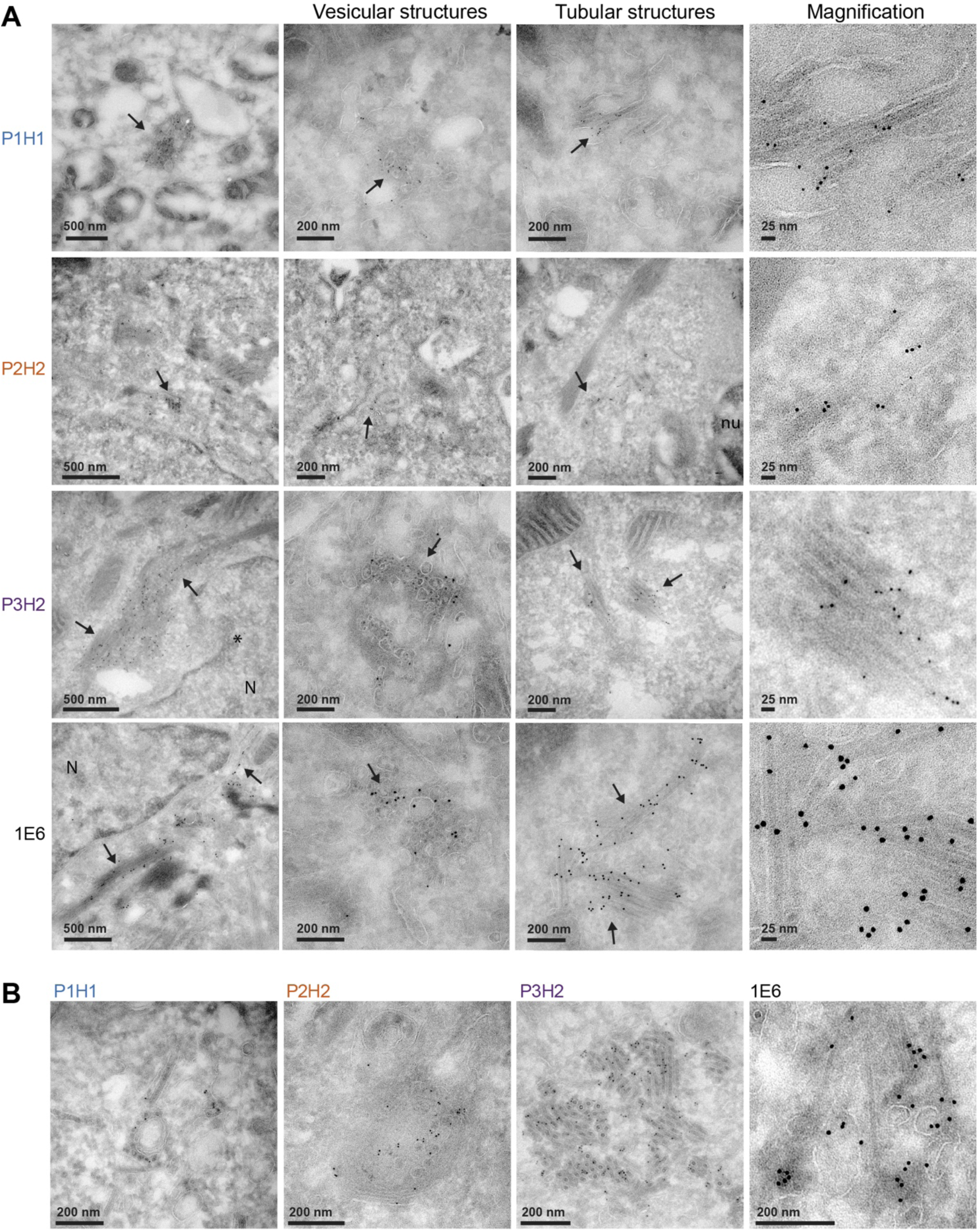
Identification of HEV-induced vesicular and tubular structures. (**A**) Transmission electron microscopy of PLC3/HEV cells cryosections immunogold- labeled with the indicated antibodies. Arrows highlight vesicular and tubular structures. N, nucleus. The asterisk indicates a nuclear deformation. (**B**) Networks containing both vesicular and tubular structures in PLC3/HEV cells.

We next performed double immunogold labeling experiments with anti-ORF2 (visualized by 6 nm gold particles) and anti-ORF3 (visualized by 10 nm gold particles) antibodies on cryosections of PLC3/HEV cells (**Fig. 4A-B**). We found a co-distribution of ORF3 and ORF2 proteins in vesicular and tubular structures, supporting the confocal microscopy analyses of ORF2/ORF3 co-localization (**Fig. 4A-B**).

**Fig 4:**
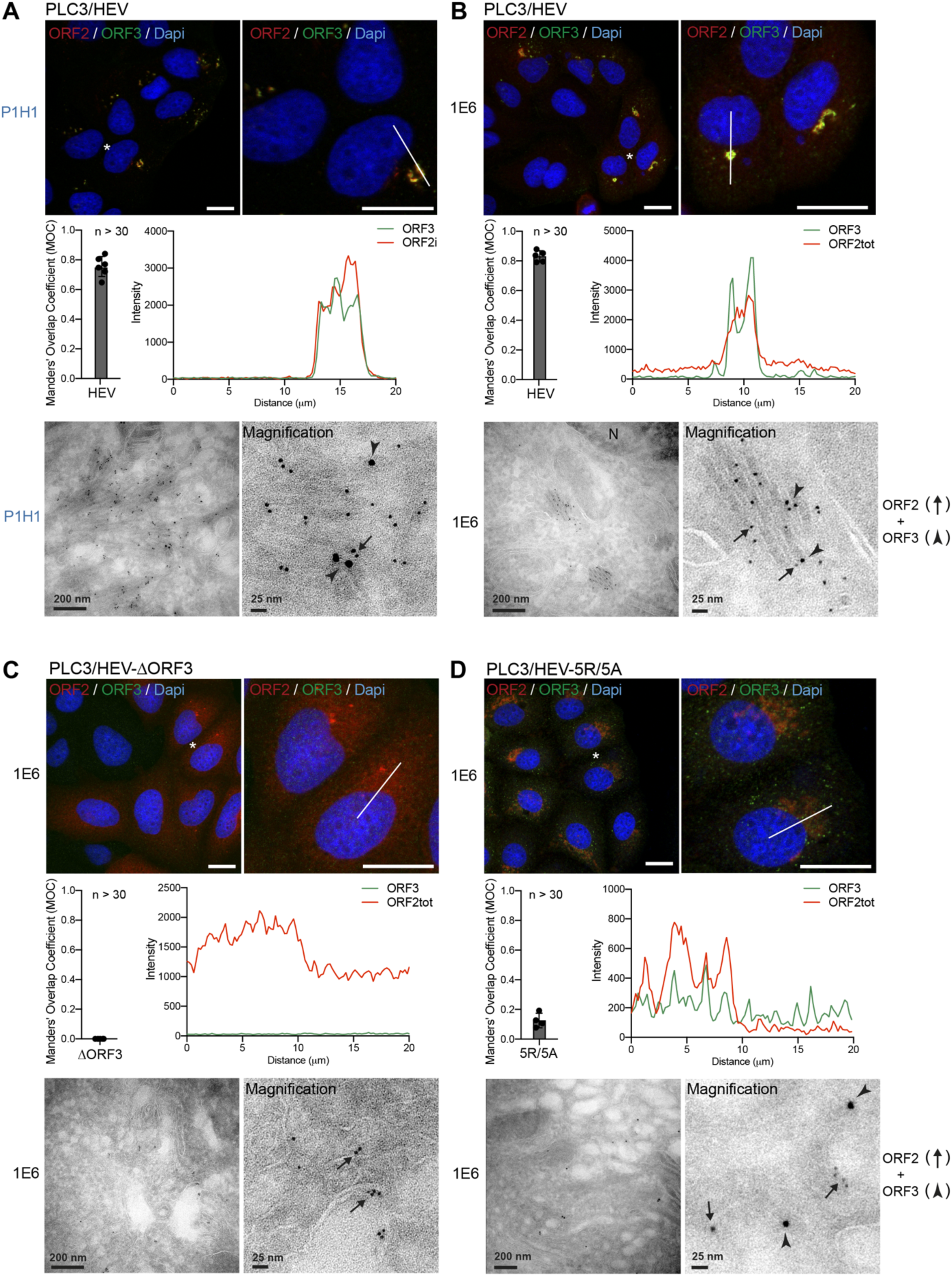
HEV-induced subcellular structures are dependent on the expression of ORF3 protein and assembly of ORF2 capsid proteins. At 6 days post- electroporation (d.p.e.), PLC3/HEV (**A, B**), PLC3/HEV-ΔORF3 (**C**) and PLC3/HEV- 5R/5A (**D**) cells were fixed, permeabilized with cold methanol and TX-0.5% and double- stained with P1H1 (**A**) or 1E6 (**B-D**) and anti-ORF3 antibodies. Red = ORF2; Green = ORF3; Blue = DAPI. Staining were analyzed by confocal microscopy. Manders’ Overlap Coefficients (MOC) of the ORF3 labeling in the ORF2 labeling were calculated on at least 30 cells. Line graphs show the fluorescence intensities of ORF2 and ORF3 staining measured every 200 nm across the region of interest highlighted by the white line in the micrograph shown above. Scale bar, 20μm. Cryosections of indicated cells were processed for double immunogold labeling with anti-ORF2 (visualized by 6 nm gold particles) and anti-ORF3 (visualized by 10 nm gold particles) antibodies, as indicated. Cryosections were next analyzed by EM. ORF2 proteins are indicated by black arrows and ORF3 proteins by arrowheads. N, nucleus. Scale bars show the indicated length.

To further understand which viral determinant is important for the formation of HEV-induced subcellular structures, we transfected PLC3 cells with an ORF3-null mutant (HEV-ΔORF3) [16] (**Fig. 4C**) and an ORF2 assembly mutant (HEV-5R/5A) [8] (**Fig. 4D**). In the absence of ORF3 protein expression, HEV particle secretion is abolished [16] and ORF2 proteins mainly accumulate in the cytosol (**Fig. 4C**). In the HEV-5R/5A mutant, the arginine residues of the ARM located in the P1 epitope (**Fig. 1A**) were replaced by alanine residues and thus prevent recognition by the P1H1 antibody. The 5R/5A mutations lead to an accumulation of ORF2 in the Golgi apparatus (**Fig. 4D**) and abrogate viral assembly but do not affect ORF3 expression level [8]. Of note, HEV RNA replication is not altered in PLC3/HEV-ΔORF3 and PLC3/HEV-5R/5A cells [8]. Strikingly, in the absence of ORF3 expression (PLC3/HEV-ΔORF3), cells mainly displayed a diffuse ORF2 staining. In EM, immunogold labeling revealed no vesicular or tubular ORF2-enriched ultrastructures but rather ORF2 proteins associated with regular cellular membranes likely derived from ER/GA compartment (**Fig. 4C**). In PLC3/HEV-5R/5A cells (**Fig. 4D**), which were characterized by an accumulation of ORF2 in the Golgi and a redistribution of ORF3 protein in the cytosol, we observed no structures resembling the ORF2/ORF3-enriched ultrastructures as those observed in **Fig. 4A-B**, but both proteins were distributed in the cytosol close to common intracellular vesicles.

Together, these results indicate that, during its lifecycle, HEV induces the formation of subcellular structures that are likely dependent on a fine interplay between ORF2 and ORF3 proteins. Of note, we did not observe any structures resembling the ORF2/ORF3-enriched ultrastructures in cells transfected with a tagged-ORF1 replicon (data not shown), indicating that these structures are not induced by HEV replication.

### HEV hijacks the Endocytic Recycling Compartment (ERC)

To further characterize the HEV-induced subcellular structures, we next carried out an extensive immunofluorescence colocalization study of the ORF2i protein with cell pathway markers. Colocalizations were performed with the P1H1 anti-ORF2i antibody and antibodies directed against markers of cytoskeleton (β-tubulin and MTOC), secretory pathway (Calnexin and ERGIC 53), early endosomes (EEA1 and Rab5), late endosomes / multivesicular bodies (MVB) (Rab9a, CD81 and CD63), Endocytic Recycling Compartment (ERC) (Rab11, CD71, EHD1, MICAL-L1 and PACSIN2), peroxisome (PMP70 and Catalase), mitochondria (TOM-20) and lysosome (LAMP1) (**Fig. 5A** and **Fig. S5**). Colocalizations were quantitatively analyzed by calculating the MOC (**Fig. 5A**). As shown in **Fig. 5A** and **5B**, the ORF2i protein significantly co-distributes with markers of two different cell compartments i.e. the late endosomes and the ERC. Indeed, the MOCs of ORF2i with Rab9a, CD81 and CD63 were 0.44, 0.55 and 0.37, respectively, indicating a medium colocalization between ORF2i and these cellular markers. Rab9a belongs to a class of small Rab GTPases which are effector proteins promoting exchanges between the late endosome pathway and the trans Golgi network [23, 24], whereas CD81 and CD63 are tetraspanins found in MVB, which are a compartment belonging to the late endosome pathway.

**Fig 5:**
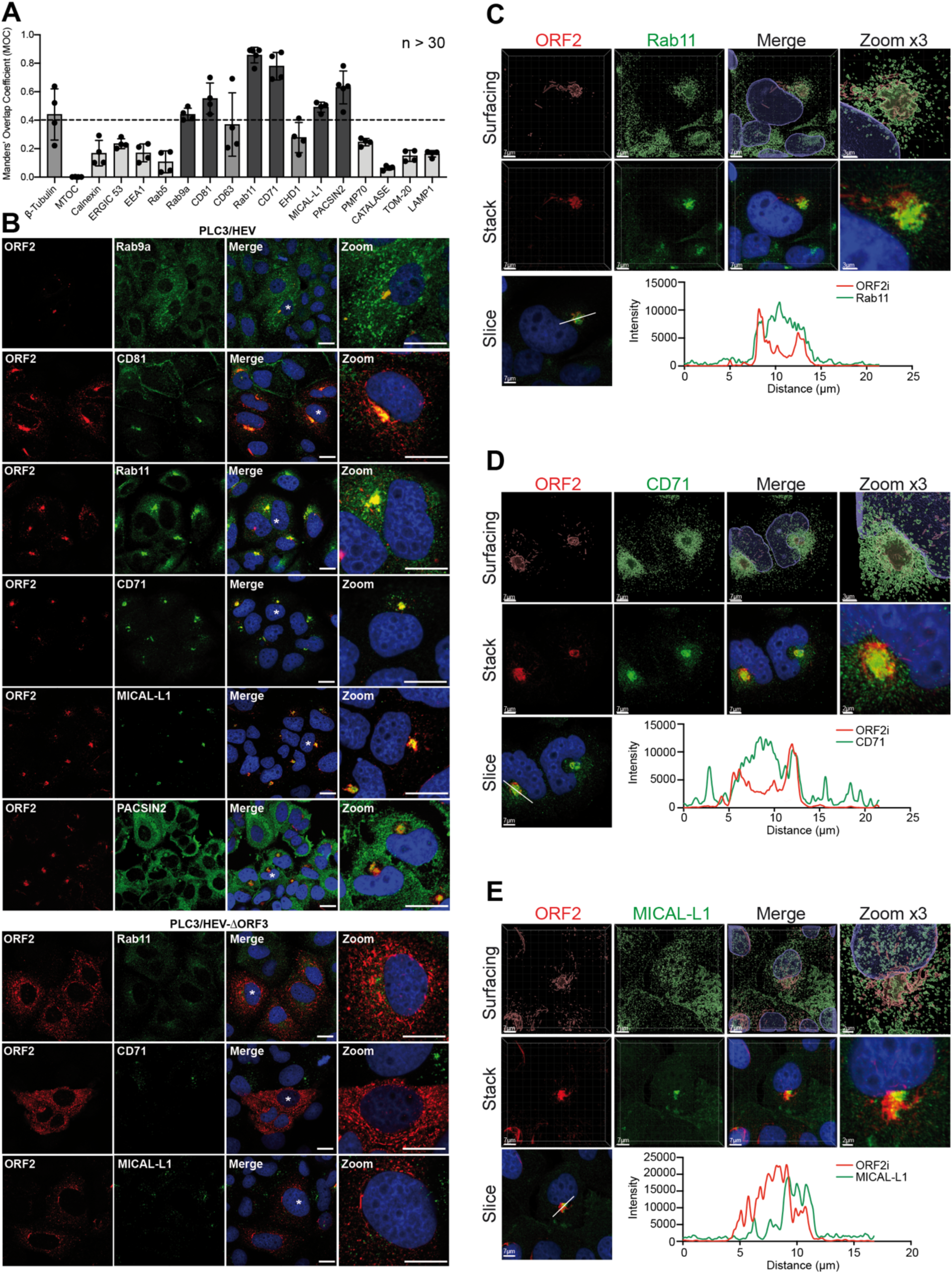
Colocalization analysis of the ORF2i protein with different cell markers. PLC3/HEV and PLC3/HEV-ΔORF3 cells were fixed, permeabilized with methanol and TX-0.5% and double-stained with P1H1 and anti-cell markers antibodies, as indicated. (**A**) Manders’ Overlap Coefficients (MOC) of ORF2 and cell marker labeling (*n* > 30 cells). Co-staining showing a low MOC are in light grey and those showing a medium MOC are in middle grey. Co-staining of PLC3/HEV cells showing a MOC > 0.4 are in dark grey and representative confocal images are shown in (**B, Top**). The co-staining of PLC3/HEV-ΔORF3 cells with P1H1 and antibodies directed against markers of recycling compartment are also shown in (**B, bottom**). Staining were analyzed by confocal microscopy. Scale bar, 20μm. PLC3/HEV cells double-stained with P1H1 and anti-Rab11 (**C**), anti-CD71 (**D**) or anti-MICAL-L1 (**E**) antibodies were next analyzed by confocal microscopy with a high resolution Airyscan module. On the top, volume rendering of the 3D z-stacks (Surfacing) using Imaris are shown to better visualize the stained substructures. In the middle, z-stacks are shown. On the bottom, line graphs show the fluorescence intensities of ORF2i and Rab11/CD71/MICAL-L1 staining measured every 50 nm across the region of interest highlighted by the white line in the micrograph shown on the bottom left of each panel. Scale bars show the indicated length. Red = ORF2; Green = cell marker; Blue = DAPI.

On the other hand, ORF2i strongly colocalized with cell markers of the ERC. This compartment is the keystone of the slow cellular recycling pathway. The ERC plays major roles in cellular metabolism and is subverted during infection by many pathogens such as viruses [25, 26]. The ERC constitutes a collection of tubular organelles that are close to the nucleus and is defined by the presence of the Rab11 GTPase and its effectors. Rab11 regulates recycling from the ERC and transport of cargo from the TGN to the plasma membrane [27–30]. Strikingly, ORF2i and Rab11 showed a MOC of 0.86 (**Fig. 5A**), indicating a strong colocalization. Indeed, super-resolution confocal microscopy analyses showed a total overlap of fluorescence intensities between ORF2i and Rab11 (**Fig. 5C**). ORF2i also strongly colocalized with CD71 (MOC=0.78) (**Fig. 5A**), the transferrin receptor that is a reference marker for the ERC [31]. This observation was further confirmed by high-resolution microscopy (**Fig. 5D**).

Efficient recycling *via* the ERC relies on the integrity of a complex network of elongated, non-symmetrical endosomes known as tubular recycling endosomes (TRE). A family of proteins known as the C-terminal Eps15 homology domain (EHD1-4) proteins and EHD-interaction partners such as MICAL-L1 (Molecule Interacting with CasL-like1) and PACSIN2/Syndapin2, are involved in TRE biogenesis and control membrane recycling. Although ORF2i only displayed a weak co-localization with EHD1 (MOC=0.28), it colocalized with MICAL-L1 (MOC=0.49) and PACSIN2 (MOC=0.63) (**Fig. 5A**). The colocalization of ORF2i with MICAL-L1 was further confirmed by high- resolution microscopy but showed a small shift of fluorescence intensities (**Fig. 5E**), indicating that they are in close proximity to each other. Altogether, our data suggest that HEV likely subverts effectors of the cellular recycling machinery.

On the other hand, although ORF2i did not colocalize with the MTOC (**Fig. 5A**), ORF2i- enriched structures were found in close proximity of the organizing center (**Fig. S5**). Of note, it has been shown that the MTOC and the ERC are two distinct structural entities closely related promoting endosomal trafficking [32].

As shown for Rab11, CD71 and MICAL-L1, colocalization analyses in PLC3/HEV- ΔORF3 cells, indicated that, in the absence of ORF3 expression, ORF2i no longer colocalizes with these cell markers (**Fig. 5B**). These results are in line with our above described observations on the importance of the ORF3 protein in the formation of HEV- induced subcellular structures (**Fig. 4**). Although some cellular markers were enriched in the HEV-induced subcellular structures (i.e. CD81 and Rab11), staining of mock electroporated PLC3 cells (PLC3 mock) (**Fig. S6**) showed similar subcellular localizations of other cell markers as in PLC3/HEV and PLC3/HEV-ΔORF3 cells (**Fig. 5**), indicating that HEV infection does not induce a general cell marker redistribution. To strengthen our previous observations, we next performed a kinetics of colocalization with the P1H1 anti-ORF2i antibody and fluorochrome-conjugated transferrin in PLC3/HEV and PLC3/HEV-ΔORF3 cells (**Fig. S7**). It has been shown that Transferrin (TrF) first binds to its receptor (CD71) at the cell membrane and then enters the cell through clathrin-mediated endocytosis. Once in the early endosomes, TrF-CD71 complexes return back to the cell surface through either a fast route going directly back to the plasma membrane, or a slower route delivering TrF-CD71 complexes to the ERC before they are trafficked back to the cell surface [31]. In PLC3/HEV cells, the colocalization between transferrin and ORF2i readily increased over time and reached a MOC of 0.70 after 45 min (**Fig. S7A**), a value similar to that found in **Fig. 5A** for its receptor. In contrast, in PLC3/HEV-ΔORF3 cells, transferrin and ORF2i showed a reduced colocalization (MOC = 0.4), (**Fig. S7B**). Thus, during the HEV lifecycle, the ORF2i protein with the help of ORF3 protein, associates with a functional ERC compartment.

We carried out double immunogold labeling experiments by combining anti-ORF2 (visualized by 10 nm gold particles) with anti-Rab11 or anti-CD71 (visualized by 6 nm gold particles) antibodies, and anti-ORF3 (10 nm) with anti-CD81 (6 nm) antibodies on cryosections of PLC3/HEV (**Fig. 6**) and PLC3 mock (**Fig. S8**) cells. In PLC3/HEV cells, we found a co-distribution of ORF2 with Rab11 and CD71 and a co-distribution of ORF3 with CD81 in vesicular and tubular structures, supporting our confocal microscopy analyses. These results indicate that the HEV-induced vesicular and tubular structures likely derive from ERC and TRE compartments.

**Fig 6:**
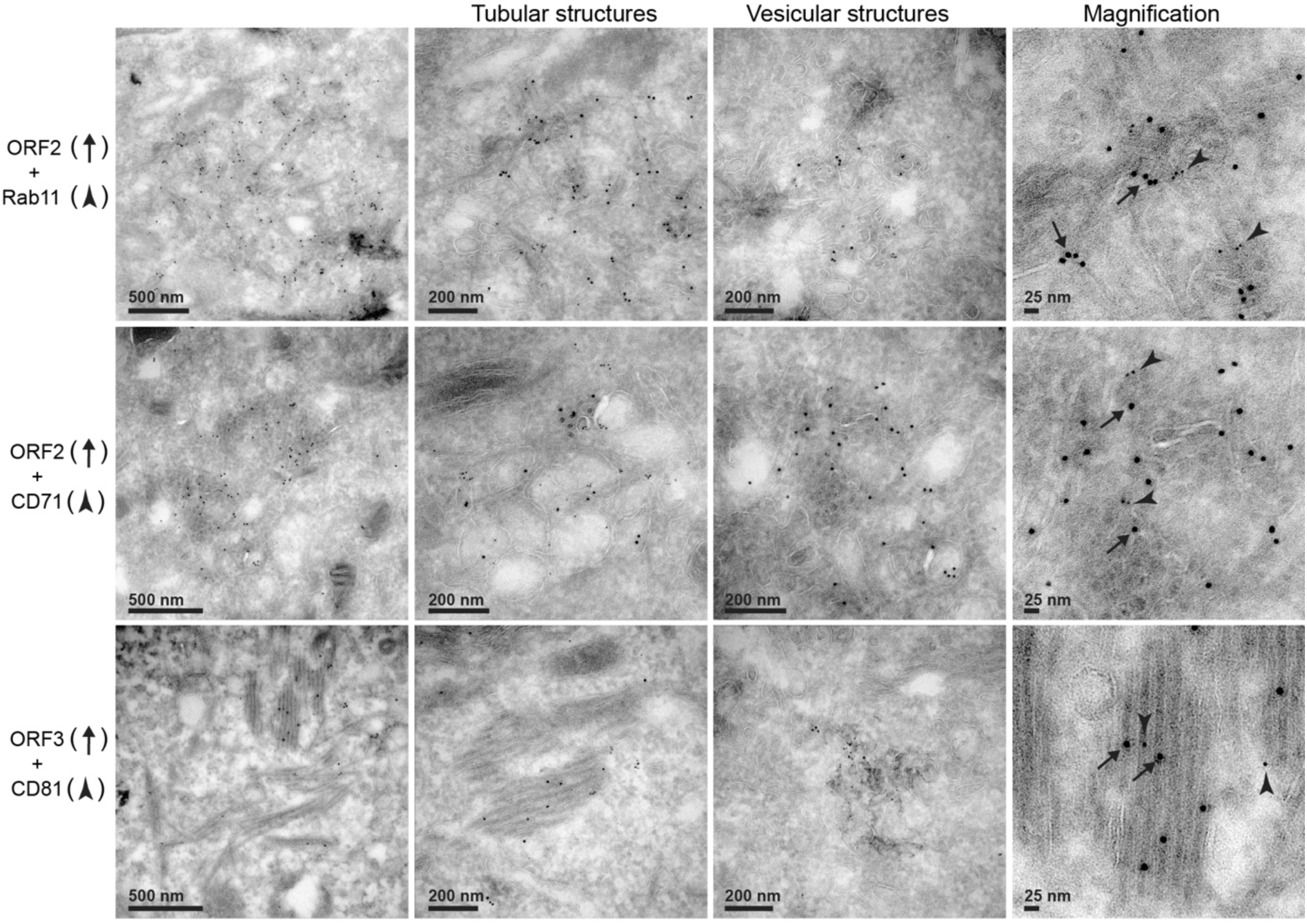
EM analysis of the co-distribution of ORF2 and ORF3 proteins with cell markers. Cryosections of PLC3/HEV cells were processed for double immunogold labeling with anti-ORF2 or anti-ORF3 (visualized by 10 nm gold particles) and anti- Rab11, anti-CD71 or anti-CD81 (visualized by 6 nm gold particles) antibodies, as indicated. Cryosections were next analyzed by EM. ORF2 and ORF3 proteins are indicated by black arrows and cell markers by arrowheads.

Moreover, the detection of CD71 and the tetraspanin CD81 confirms the presence of membranes in vesicular and tubular structures. Since ORF3 protein is a key player in the biogenesis of eHEV [33] and CD81 is present on the quasi-envelope [34], the ORF2/ORF3-enriched ultrastructures that we identified likely correspond to eHEV viral factories. In line with this, we recently demonstrated that the ORF1 replicase and genomic RNA are co-distributed with ORF2, ORF3 and Rab11 proteins in nugget-like structures (**Fig S9** and [17]). In addition, P1H1, P2H2, P3H2 and 1E6 anti-ORF2 immunolabeling on cryosections of PLC3/HEV cells highlighted in the ORF2-enriched membranous compartments numerous viral-like particles of ∼25 nm in diameter (**Fig. 7**).

**Fig 7:**
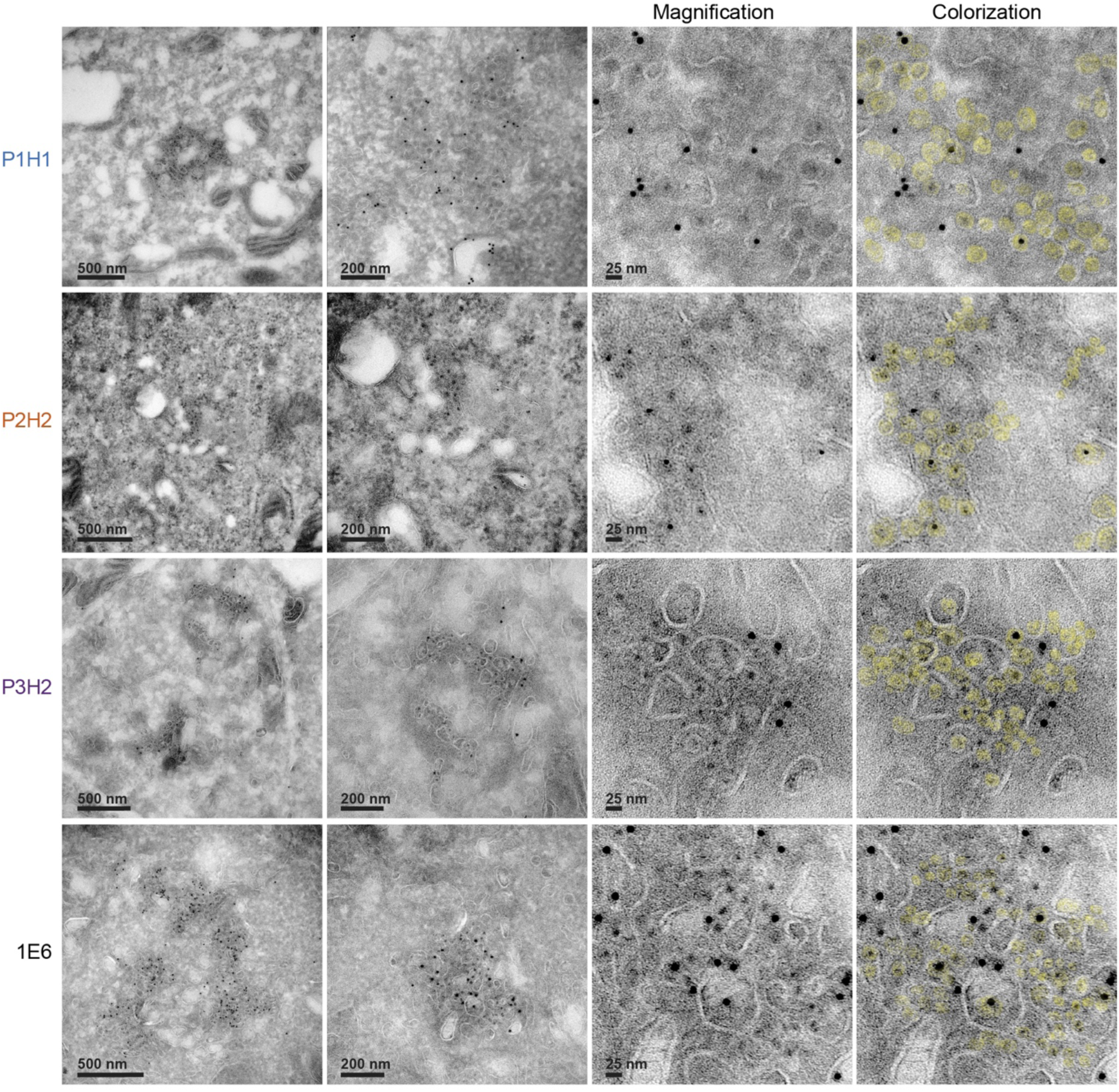
Visualization of intracellular HEV like-particles. Cryosections of PLC3/HEV cells were immunogold-labeled with the indicated antibodies and analyzed by EM. On the right column, particles were highlighted by colorization in yellow.

### The ERC plays a central role in the HEV lifecycle

To confirm that the ERC plays a central role in the HEV lifecycle, we conducted functional studies. PLC3/HEV cells were transfected with small interfering RNA (siRNA) targeting Rab11a and Rab11b isoforms (siRab11) or non-targeting siRNA (siCTL) (**Fig. 8**). The effect of Rab11 silencing on intracellular ORF2 expression was analyzed by IF (**Fig. 8A**) and WB (**Fig. 8B**). The effect of Rab11 silencing on viral production was analyzed by quantification of secreted viral RNA (**Fig. 8C**) and infectious titers (**Fig. 8D**). The effect of Rab11 silencing on genome replication was analyzed by using PLC3 cells stably harboring a subgenomic replicon (**Fig. 8E**).

**Fig 8:**
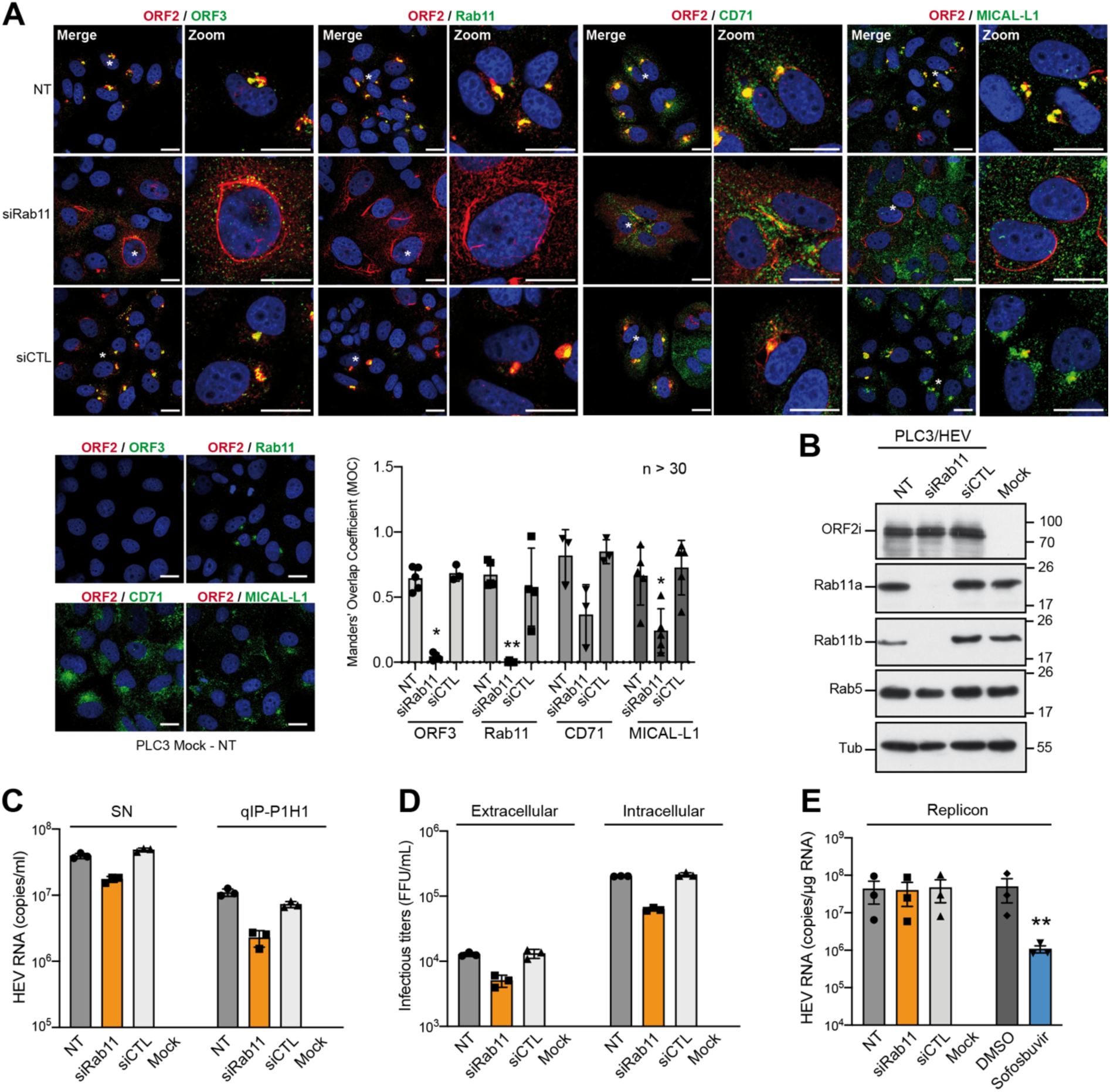
Effect of Rab11 silencing on protein expression and HEV particle secretion. (**A-D**) PLC3/HEV cells were transfected with siRNA targeting Rab11a and Rab11b (siRab11), with a non-targeting siRNA control (siCTL) or left non-transfected (NT). Non-electroporated PLC3 cells were used as controls (Mock). At 72h post- transfection, cells were analyzed by IF (**A**) and WB (**B**) using the indicated antibodies. Staining were analyzed by confocal microscopy. Scale bar, 20μm. Red = ORF2 stained by P1H1; Green = ORF3 or cell marker; Blue = DAPI. Manders’ Overlap Coefficients were calculated. (**C**) Quantification of HEV RNA in SN of transfected cells was performed by direct RT-qPCR (SN) and after IP with the P1H1 antibody (qIP-P1H1). (**D**) Production of extracellular and intracellular infectious particles in transfected cells was evaluated by viral titration. (**E**) PLC3 cells stably replicating a p6 subgenomic replicon were transfected with siRNAs as described above. At 72h post-transfection, intracellular HEV RNA were quantified by RT-qPCR. Mock cells were used as a negative control. PLC3 cells harboring the replicon and treated with 20μM of Sofosbuvir were used as a positive control for replication inhibition.

Although Rab11 silencing did not affect the ORF2 expression level, we found that it abrogated the formation of nugget-like structures to the benefit of ORF2-enriched stringy structures mostly localized around the nucleus. These stringy structures did not co-distribute with ORF3, CD71 nor MICAL-L1, which display a more diffuse pattern upon Rab11 silencing (**Fig. 8A**). Moreover, extracellular RNA levels and infectious titers were reduced in siRab11-transfected PLC3/HEV cells, as compared to control cells (NT and siCTL) (**Fig. 8C-D**). Although no effect of siRab11 on HEV replication was observed (**Fig. 8E)**, infectious titers showed a significant reduction (i.e., 70%) in intracellular progeny (**Fig. 8D)**, indicating that disruption of the ERC by Rab11 silencing impairs HEV particle assembly. Altogether, these results confirm that the hijacking of the recycling compartment by HEV is pivotal for producing viral particles.

## Discussion

The study performed here with the help of home-made anti-ORF2 antibodies, notably antibodies recognizing the particle associated-ORF2i form, and HEV-producing cells brings new insights into the HEV-host cell interactions.

Previously, we and other demonstrated that during HEV infection, different isoforms of the ORF2 capsid protein are secreted [6, 7]. Indeed, by combining the gt3 p6 strain [15] and the highly transfectable PLC3 cells, we identified 3 forms of the ORF2 capsid protein that perform distinct functions in the HEV lifecycle and display different sequences, post-translational modifications and subcellular localization [6, 14]. The ORF2i form is the component of infectious particles and derives from the assembly of intracellular ORF2i protein [8]. The ORF2i protein is not glycosylated and its sequence starts at Leu14 corresponding to the middle of the signal peptide (Fig. 1A). Although it has been shown that ORF2i protein might be produced from an additional start codon [7], we recently found that an Arginine-Rich Motif (ARM) located in the ORF2 N-terminal region regulates the dual topology and functionality of ORF2 signal peptide, leading to the production of either the cytosolic infectious ORF2i that is not processed by the signal peptidase or the ORF2g/c forms that are generated by translocation into the ER lumen [8]. The glycosylated ORF2g/c forms, which are not associated with infectious virions and likely act as immunological baits [6, 7], are produced by translocation of ORF2 proteins into the secretory pathway where they are highly glycosylated, cleaved by furin, and quickly secreted [6, 7]. Here, we capitalized on these distinctive features to generate and characterize four anti-ORF2 monoclonal antibodies, including three antibodies (P1H1, P2H1 and P2H2) directed against the ORF2i form and one antibody (P3H2) directed against the different ORF2 isoforms (Fig. 1). Analyses by WB and IP of HEV-infected cell lysates showed that the four antibodies equally recognize the intracellular ORF2 proteins. In contrast, analyses on HEV-infected cell supernatants, which contain some ORF2i proteins but also huge amounts of ORF2g/c proteins, demonstrated that the P1H1, P2H1 and P2H2 antibodies specifically recognize the ORF2i protein without cross-reaction with the glycosylated ORF2 forms. Importantly, we found that the P1H1 antibody recognizes non-lipid-associated HEV particles from cell culture and patient sera. Therefore, the P1H1 antibody might represent a good candidate for diagnosis purposes. More generally, the antibodies that we generated in this study represent unique tools for deciphering the biogenesis mechanisms of ORF2 isoforms and their precise functions in the HEV lifecycle.

Structure of the ORF2 N-terminal domain has not been resolved yet. It is not included in most of the recombinant constructs that have yielded structural data, and it is disordered in the single structure of full-size particle available [18]. However, based on its potential interaction with RNA and its N-terminal location to the innermost S (shell) domain, this N-terminal domain is thought to be orientated toward the inner cavity of particles. In our study, we found that the P1H1 antibody that recognizes the N-terminal domain of ORF2 (Fig. 1A) efficiently captures delipidated HEV particles (Fig. 1D-F). Although we cannot exclude that the P1H1 antibody recognizes partially assembled ORF2i proteins, our results suggest that the P1 epitope might be exposed on the viral surface. Our structural models showed that this is readily possible (Fig. 1G and 1H): At least some N-termini may reach as high as or higher than the outer P (protruding) domain and display the P1 epitope, while keeping most other arginine residues directed toward the capsid interior. In addition, we recently found that the ARM located in the ORF2 N-terminal region promotes ORF2 membrane association that is likely essential for particle assembly [8]. Therefore, we hypothesize that the ORF2 N-terminus is associated with eHEV-enveloping lipids and removal of lipids by detergent treatment unmasks ORF2 N-terminal epitopes, including the P1 epitope.

In the present study, we identified HEV-induced subcellular structures that are enriched in viral proteins. Indeed, we found that ORF2 and ORF3 proteins highly co- localize in HEV-producing and infected cells. In addition, thanks to the P1H1 antibody that specifically recognizes the particle-associated ORF2i protein, we demonstrated that ORF2i and ORF3 proteins are tightly co-distributed in perinuclear nugget-like structures, as observed in confocal microscopy. In the absence of ORF3 expression, we observed a redistribution of the ORF2i protein in cytosolic dot-like structures, and in the absence of ORF2 assembly, the ORF3 protein was as well redistributed through the cytosol, indicating that both proteins are tightly connected. Ultrastructural analyses by EM of cryosections of HEV-replicating cells showed that the ORF2/ORF3 enriched structures correspond to a network of vesicular and tubular components located in the vicinity of the nucleus. While vesicular structures were heterogeneous in their organization and their size i.e. 50-250 nm, the tubular structures formed regular parallel arrays and displayed a homogeneous diameter of 20-25nm, corresponding to that of intracellular HEV-like particles observed in cryosections. Therefore, we hypothesize that tubular structures might correspond to virion precursors containing assembled or pre-assembled virions. Vesicular and tubular structures were found in both PLC3/HEV and Huh-7.5/HEV cells, indicating that they are not cell type-specific. Moreover, we found the same structures in HEV-infected Huh-7.5 cells indicating that they do not correspond to an artefact of cell electroporation. Although difficult to set up, it would be also interesting to analyze whether this kind of structures is formed in heterologous expression system and in other infectious models such as cells replicating gt1 or in HEV-infected primary cells.

Interestingly, in the absence of ORF2 assembly or ORF3 expression, no perinuclear nugget-like shapes were observed in confocal microscopy, nor vesicular and tubular structures in EM. ORF3 protein is not involved in virion assembly but plays an essential role in exosomal release and acquisition of the quasi-envelope around the neo- synthesized viral particles [35]. In addition, recent reports by Szkolnicka *et al.* and our laboratory who studied the subcellular localization of the ORF1 replicase using epitope-tagged replicons and full-length genomes found replication complexes in cytoplasmic dot-like structures overlapping significantly with ORF2 and ORF3 proteins as well as HEV RNA ([17, 36] and Fig S9). We can therefore speculate that the observed structures represent viral factories in which assembled particles acquire their membrane through an ORF2-ORF3 interaction.

The ORF2 subcellular localization has been studied in heterologous systems and found in different cell compartments i.e. plasma membrane, ER and cytoplasmic compartment [37, 38]. Later on, ORF2 characterization in infectious systems led to the identification of the ORF2 isoforms which are found partitioned in different subcellular compartments. The glycosylated ORF2 forms go through the secretory pathway. The ORF2i form is mainly present in the cytosolic compartment but is also translocated into the nucleus of infected cells [8,14,39]. Here, thanks to the specificity of our antibodies, notably the P1H1 antibody, we performed an extensive colocalization study of ORF2i proteins with different cell markers. Of note, as immunofluorescence analysis of ORF2 expression was performed at 6 days p.e., the nuclear localization of the protein could not be shown here as nuclear translocation was reported to occur at early time points p.e. (i.e. 18 hours p.e.) [8, 14]. In accordance with previous studies [20, 40], we found a partial co-localization of ORF2i with late endosome markers notably with the tetraspanin CD81, a marker of MVB and exosomes. Since ORF1 and ORF3 proteins have been also shown to colocalize with MVB markers [20, 36], our findings strengthen the hypothesis of a close connection between HEV replication and assembly sites.

A recent study showed that ORF3 is palmitoylated at cysteine residues in its N-terminal region and is exposed to the cytosolic side of the intracellular membranes [22]. It has also been localized in early and recycling endosomes [41] as well as in MVB [20] and has been found in association with microtubules [42, 43]. Hence, ORF3 may behave as a cargo able to drive ORF2 to HEV replication/assembly site.

Importantly, we identified a strong colocalization of ORF2i and ORF3 with several markers of the recycling compartment, which was abrogated upon Rab11 silencing. ERC is involved in several stages of the lifecycle of a number of DNA and RNA viruses, with Rab11 being a central player in most of these processes [26]. ERC notably mediates viral transport, assembly and egress e.g. it is involved in the envelopment of herpes simplex-1 capsids [44] or contributes to the transport of HCV virions towards the plasma membrane [45]. Here, we demonstrated that HEV particle assembly depends on a functional ERC. Interestingly, it has been shown that ERC hijacking is associated with membrane remodeling upon infection. Cholesterol accumulates at the ERC during Influenza A virus (IAV) infection, or Rab11 redistributes from dot structures to large aggregates during infection with several viruses including IAV [26]. Ultrastructural changes of these ERC membrane remodeling were poorly investigated to date. In our study, we demonstrated that HEV is a new candidate in the list of viruses hijacking the ERC. Importantly, we found that viral proteins and recycling compartment markers are co-distributed in perinuclear structures found in ultrastructural analyses as a network of vesicular and tubular structures. To our knowledge, this kind of structures has never been described before and might be the hallmark of HEV infection.

## ACKNOWLEDGMENTS

This work was supported by the French agency ANRS-Maladies infectieuses émergentes. This work was also supported by the Pasteur Institute of Lille, Région Hauts-de-France, Inserm-transfert and the University of Tours. C.B. and K.H. were supported by a fellowship from the ANRS. C.C. and M.F. were supported by the Pasteur Institute of Lille and Région Hauts-de-France. C.C. was also supported by Inserm-transfert. We thank Suzanne U. Emerson (NIH, USA), Jérôme Gouttenoire (University of Lausanne) and Ralf Bartenschlager (University of Heidelberg) for providing us with reagents. We thank Valentin de Masson d’Autume for his technical contribution. We thank Raphaël Guérois for sharing a standalone ColabFold implementation of AlphaFold2 and the I2BC integrative bioinformatics core facility BIOI2 for assistance with the high-performance computing infrastructure.

## Abbreviations

aa: amino acid
ARM: Arginine-Rich Motif
EEA1: Early Endosome Antigen-1
EHD: Eps15 homology domain
eHEV: enveloped HEV
EM: Electron Microscopy
ER: Endoplasmic Reticulum
ERC: Endocytic-Recycling Compartment
ERGIC: Endoplasmic Reticulum-Golgi Intermediate Compartment
ESCRT: Endosomal Sorting Complexes Required for Transport
GA: Golgi apparatus
gt: genotype
HD: Heat-denatured
HEV: Hepatitis E Virus
IAV: Influenza A virus
IF: Immunofluorescence
IG: Immunogold
IP: Immunoprecipitation
MICAL-L1: Molecule Interacting with CasL-like1
MOC: Mander’s Overlap Coefficient
MTOC: Microtubule-Organizing Center
MVB: Multivesicular Bodies
ORF: Open Reading Frame
ORF2c: cleaved ORF2
ORF2g: glycosylated ORF2
ORF2i: infectious ORF2
PACSIN2: Protein Kinase C and Casein Kinase Substrate in Neurons 2
p.e.: post-electroporation
PMP70: 70-kDa Peroxisomal Membrane Protein
qIP: immunoprecipitation followed by RT-qPCR
siRNA: small interfering RNA
SN: Supernatant
TRE: Tubular-Recycling Endosome
TrF: Transferrin
TX: Triton X-100
WB: Western Blotting

## References

1. Kamar N, Izopet J, Pavio N, Aggarwal R, Labrique A, Wedemeyer H, et al. Hepatitis E virus infection. Nature Reviews Disease Primers 2017;3:17086. https://doi.org/10.1111/trf.13355.

2. Doceul V, Bagdassarian E, Demange A, Pavio N. Zoonotic Hepatitis E Virus: Classification, Animal Reservoirs and Transmission Routes. Viruses 2016;8:270. https://doi.org/10.3390/v8100270.

3. Horvatits T, Wiesch JS zur, Lütgehetmann M, Lohse AW, Pischke S. The Clinical Perspective on Hepatitis E. Viruses 2019;11:617–9. https://doi.org/10.3390/v11070617.

4. Aslan AT, Balaban HY. Hepatitis E virus: Epidemiology, diagnosis, clinical manifestations, and treatment. World J Gastroentero 2020;26:5543–60. https://doi.org/10.3748/wjg.v26.i37.5543.

5. Nimgaonkar I, Ding Q, Schwartz RE, Ploss A. Hepatitis E virus: advances and challenges. Nature Reviews Gastroenterology & Hepatology 2018;15:96–110. https://doi.org/10.1038/nrgastro.2017.150.

6. Montpellier C, Wychowski C, Sayed IM, Meunier J-C, Saliou J-M, Ankavay M, et al. Hepatitis E Virus Lifecycle and Identification of 3 Forms of the ORF2 Capsid Protein. Gastroenterology 2018;154:211–223.e8. https://doi.org/10.1053/j.gastro.2017.09.020.

7. Yin X, Ying D, Lhomme S, Tang Z, Walker CM, Xia N, et al. Origin, antigenicity, and function of a secreted form of ORF2 in hepatitis E virus infection. Proceedings of the National Academy of Sciences of the United States of America 2018;3:201721345–6. https://doi.org/10.1073/pnas.1721345115.

8. Hervouet K, Ferrié M, Ankavay M, Montpellier C, Camuzet C, Alexandre V, et al. An Arginine-Rich Motif in the ORF2 Capsid Protein Regulates the Hepatitis E Virus Lifecycle and Interactions with the Host Cell. Biorxiv 2021:2021.05.26.445820. https://doi.org/10.1101/2021.05.26.445820.

9. Meister TL, Bruening J, Todt D, Steinmann E. Cell culture systems for the study of hepatitis E virus. Antiviral Research 2019;163:34–49. https://doi.org/10.1016/j.antiviral.2019.01.007.

10. Ju X, Ding Q. Hepatitis E Virus Assembly and Release. Viruses 2019;11:539–13. https://doi.org/10.3390/v11060539.

11. Harak C, Lohmann V. Ultrastructure of the replication sites of positive-strand RNA viruses. Virology 2015;479:418–33. https://doi.org/10.1016/j.virol.2015.02.029.

12. Talmont F, Moulédous L, Baranger M, Gomez-Brouchet A, Zajac J-M, Deffaud C, et al. Development and characterization of sphingosine 1-phosphate receptor 1 monoclonal antibody suitable for cell imaging and biochemical studies of endogenous receptors. PLoS ONE 2019;14:e0213203–19. https://doi.org/10.1371/journal.pone.0213203.

13. Blight KJ, Mckeating JA, Rice CM. Highly Permissive Cell Lines for Subgenomic and Genomic Hepatitis C Virus RNA Replication. Journal of Virology 2002;76:13001–14. https://doi.org/10.1128/jvi.76.24.13001-13014.2002.

14. Ankavay M, Montpellier C, Sayed IM, Saliou J-M, Wychowski C, Saas L, et al. New insights into the ORF2 capsid protein, a key player of the hepatitis E virus lifecycle. Scientific Reports 2019;9:6243. https://doi.org/10.1038/s41598-019-42737-2.

15. Shukla P, Nguyen HT, Faulk K, Mather K, Torian U, Engle RE, et al. Adaptation of a genotype 3 hepatitis E virus to efficient growth in cell culture depends on an inserted human gene segment acquired by recombination. Journal of Virology 2012;86:5697–707. https://doi.org/10.1128/jvi.00146-12.

16. Graff J, Nguyen H, Yu C, Elkins WR, Claire MS, Purcell RH, et al. The Open Reading Frame 3 Gene of Hepatitis E Virus Contains a cis-Reactive Element and Encodes a Protein Required for Infection of Macaques. Journal of Virology 2005;79:6680–9. https://doi.org/10.1128/jvi.79.11.6680-6689.2005.

17. Metzger K, Bentaleb C, Hervouet K, Alexandre V, Montpellier C, Saliou J-M, et al. Processing and Subcellular Localization of the Hepatitis E Virus Replicase: Identification of Candidate Viral Factories. Front Microbiol 2022;13:828636. https://doi.org/10.3389/fmicb.2022.828636.

18. Xing L, Li TC, Mayazaki N, Simon MN, Wall JS, Moore M, et al. Structure of hepatitis E virion-sized particle reveals an RNA-dependent viral assembly pathway. Journal of Biological Chemistry 2010;285:33175–83. https://doi.org/10.1074/jbc.m110.106336.

19. Ding Q, Heller B, Capuccino JMV, Song B, Nimgaonkar I, Hrebikova G, et al. Hepatitis E virus ORF3 is a functional ion channel required for release of infectious particles. PNAS 2017:201614955. https://doi.org/10.1073/pnas.1614955114.

20. Nagashima S, Jirintai S, Takahashi M, Kobayashi T, Tanggis, Nishizawa T, et al. Hepatitis E virus egress depends on the exosomal pathway, with secretory exosomes derived from multivesicular bodies. Journal of General Virology 2014;95:2166–75. https://doi.org/10.1099/vir.0.066910-0.

21. Nagashima S, Takahashi M, Jirintai, Tanaka T, Yamada K, Nishizawa T, et al. A PSAP motif in the ORF3 protein of hepatitis E virus is necessary for virion release from infected cells. Journal of General Virology 2011;92:269–78. https://doi.org/10.1099/vir.0.025791-0.

22. Gouttenoire J, Pollán A, Abrami L, Oechslin N, Mauron J, Matter M, et al. Palmitoylation mediates membrane association of hepatitis E virus ORF3 protein and is required for infectious particle secretion. PLoS Pathogens 2018;14:e1007471–24. https://doi.org/10.1371/journal.ppat.1007471.

23. Lombardi D, Soldati T, Riederer MA, Goda Y, Zerial M, Pfeffer SR. Rab9 functions in transport between late endosomes and the trans Golgi network. Embo J 1993;12:677–82. https://doi.org/10.1002/j.1460-2075.1993.tb05701.x.

24. Hutagalung AH, Novick PJ. Role of Rab GTPases in Membrane Traffic and Cell Physiology. Physiol Rev 2011;91:119–49. https://doi.org/10.1152/physrev.00059.2009.

25. Maxfield FR, McGraw TE. Endocytic recycling. Nature Reviews Molecular Cell Biology 2004;5:121–32. https://doi.org/10.1038/nrm1315.

26. Vale-Costa S, Amorim M. Recycling Endosomes and Viral Infection. Viruses 2016;8:64–29. https://doi.org/10.3390/v8030064.

27. Ullrich O, Reinsch S, Urbé S, Zerial M, Parton RG. Rab11 regulates recycling through the pericentriolar recycling endosome. J Cell Biology 1996;135:913–24. https://doi.org/10.1083/jcb.135.4.913.

28. Stone R, Hayashi T, Bajimaya S, Hodges E, Takimoto T. Critical role of Rab11a- mediated recycling endosomes in the assembly of type I parainfluenza viruses. Virology 2016;487:11–8. https://doi.org/10.1016/j.virol.2015.10.008.

29. Bhuin T, Roy JK. Rab11 in disease progression. Int J Mol Cell Medicine 2015;4:1–8.

30. Guichard A, Nizet V, Bier E. RAB11-mediated trafficking in host–pathogen interactions. Nat Rev Microbiol 2014;12:624–34. https://doi.org/10.1038/nrmicro3325.

31. Mayle KM, Le AM, Kamei DT. The intracellular trafficking pathway of transferrin. Biochimica Et Biophysica Acta Bba - Gen Subj 2012;1820:264–81. https://doi.org/10.1016/j.bbagen.2011.09.009.

32. Hehnly H, Chen C-T, Powers CM, Liu H-L, Doxsey S. The Centrosome Regulates the Rab11- Dependent Recycling Endosome Pathway at Appendages of the Mother Centriole. Curr Biol 2012;22:1944–50. https://doi.org/10.1016/j.cub.2012.08.022.

33. Yang Y-L, Nan Y-C. Open reading frame 3 protein of hepatitis E virus: Multi-function protein with endless potential. World J Gastroentero 2021;27:2458–73. https://doi.org/10.3748/wjg.v27.i20.2458.

34. Nagashima S, Takahashi M, Kobayashi T, Tanggis, Nishizawa T, Nishiyama T, et al. Characterization of the Quasi-Enveloped Hepatitis E Virus Particles Released by the Cellular Exosomal Pathway. Journal of Virology 2017;91. https://doi.org/10.1128/jvi.00822-17.

35. Glitscher M, Hildt E. Hepatitis E virus egress and beyond – the manifold roles of the viral ORF3 protein. Cell Microbiol 2021:e13379. https://doi.org/10.1111/cmi.13379.

36. Szkolnicka D, Pollán A, Silva ND, Oechslin N, Gouttenoire J, Moradpour D. Recombinant Hepatitis E Viruses Harboring Tags in the ORF1 Protein. Journal of Virology 2019;93:1237–18. https://doi.org/10.1128/jvi.00459-19.

37. Surjit M, Jameel S, Lal SK. Cytoplasmic localization of the ORF2 protein of hepatitis E virus is dependent on its ability to undergo retrotranslocation from the endoplasmic reticulum. Journal of Virology 2007;81:3339–45. https://doi.org/10.1128/jvi.02039-06.

38. Zafrullah M, Ozdener MH, Kumar R, Panda SK, Jameel S. Mutational analysis of glycosylation, membrane translocation, and cell surface expression of the hepatitis E virus ORF2 protein. Journal of Virology 1999;73:4074–82.

39. Lenggenhager D, Gouttenoire J, Malehmir M, Bawohl M, Honcharova-Biletska H, Kreutzer S, et al. Visualization of hepatitis E virus RNA and proteins in the human liver. Journal of Hepatology 2017;67:471–9. https://doi.org/10.1016/j.jhep.2017.04.002.

40. Nagashima S, Takahashi M, Jirintai S, Tanaka T, Nishizawa T, Yasuda J, et al. Tumour susceptibility gene 101 and the vacuolar protein sorting pathway are required for the release of hepatitis E virions. Journal of General Virology 2011;92:2838–48. https://doi.org/10.1099/vir.0.035378-0.

41. Chandra V, Kar-Roy A, Kumari S, Mayor S, Jameel S. The Hepatitis E Virus ORF3 Protein Modulates Epidermal Growth Factor Receptor Trafficking, STAT3 Translocation, and the Acute-Phase Response. Journal of Virology 2008;82:7100–10. https://doi.org/10.1128/jvi.00403-08.

42. Zafrullah M, Ozdener MH, Panda SK, Jameel S. The ORF3 protein of hepatitis E virus is a phosphoprotein that associates with the cytoskeleton. Journal of Virology 1997;71:9045–53.

43. Kannan H, Fan S, Patel D, Bossis I, Zhang Y-J. The hepatitis E virus open reading frame 3 product interacts with microtubules and interferes with their dynamics. Journal of Virology 2009;83:6375–82. https://doi.org/10.1128/jvi.02571-08.

44. Hollinshead M, Johns HL, Sayers CL, Gonzalez-Lopez C, Smith GL, Elliott G. Endocytic tubules regulated by Rab GTPases 5 and 11 are used for envelopment of herpes simplex virus. The EMBO Journal 2012;31:4204–20. https://doi.org/10.1038/emboj.2012.262.

45. Coller KE, Heaton NS, Berger KL, Cooper JD, Saunders JL, Randall G. Molecular Determinants and Dynamics of Hepatitis C Virus Secretion. Plos Pathog 2012;8:e1002466. https://doi.org/10.1371/journal.ppat.1002466.

