## Supplementary information text and figures for "The Endocytic Recycling Compartment Serves as a Viral Factory for Hepatitis E Virus"

**Content**


**Supplementary results**

**Supplementary Materials and methods**

- Preparation of immunogens.
- Immunization of mice.
- Cells
- Plasmids and transfection
- Modeling of the N-terminus of ORF2.
- Patient samples.
- RT-qPCR.
- Western blotting analyses.
- Indirect immunofluorescence.
- Transferrin endocytosis.
- Infectious titers
- Intracellular viral particles
- *In situ* labeling of viral RNA

**Supplementary Figure legends**

Supplementary Figure 1: ORF2 and ORF3 antibody recognition in PLC3 mock cells.

Supplementary Figure 2: Antibody recognition of gt1 and gt3 ORF2 proteins.

Supplementary Figure 3: Immunogold labeling of PLC3 and Huh-7.5 mock cells.

Supplementary Figure 4: Identification of HEV-induced vesicular and tubular structures in HEV-producing Huh-7.5 cells.

Supplementary Figure 5: Colocalization analysis of the ORF2i protein with different cell markers in PLC3/HEV cells.

Supplementary Figure 6: Colocalization analysis of the ORF2i protein with different cell markers in PLC3 mock cells.

Supplementary Figure 7: Kinetics of colocalization of the ORF2i protein with transferrin.

Supplementary Figure 8: Double-immunogold labeling of PLC3 mock cells.

Supplementary Figure 9: Detection of ORF1 protein and genomic HEV RNA in HEV-producing PLC3 cells.

**Supplementary table**

Supplementary Table 1: Primary antibodies used in Western-blot (WB), immunofluorescence (IF) and immunogold (IG) experiments.

### References

### Supplementary results

#### Antibody recognition of gt1 and gt3 ORF2 protein.

Since no efficient cell culture model of HEV gt1 is currently available, we used a heterologous expression system to compare antibody recognition towards gt1 and gt3 ORF2 proteins. Sar55 (gt1) and p6 (gt3) ORF2 sequences were cloned into pTM plasmids and expressed in Huh-7 cells stably expressing the T7 RNA-polymerase (H7-T7-IZ cells) [1]. As shown in **Fig. S2**, although the ORF2-gt3 pattern was different from that observed in infectious system, the ORF2 recognition pattern for each antibody was similar for gt1 and gt3 ORF2-expressing cells. These results indicate that antibodies recognized ORF2 proteins from HEV-gt1.

### Supplementary Materials and methods

**Preparation of immunogens.** Peptides P1 (GQPSGRRRGRRSGG), P2 (AGYPYNYNTTASDQ) and P3 (SRVVIQDYDNQHEQDR) were synthetized and coupled to the protein carrier KLH *via* a maleimide function by adding a cysteine at the C-terminal position.

**Immunization of mice.** BIOTEM animal experiments were performed in accordance with the dedicated laws. Institutional Animal Care Committee (IACUC) was DDPP de l'Isère. BIOTEM Ethics committee was the approving committee. Peptides in complete Freund's adjuvant were injected (20µg) subcutaneously in OF1 mice. Mice were subsequently immunized twice with peptides (40µg) in incomplete Freund's adjuvant. Peptides were administrated intraperitoneally in mice three days before cell fusion between splenocytes and myeloma cell line (NS-1). The selection of secreting hybridomas was performed in Hypoxanthine-Aminopterin-Thymidine medium. Before being euthanized using carbon dioxide method, mice were bled to collect immune serum. Screening of sera, hybridomas and subclones was performed by western-blotting and immunofluorescence on PLC3/HEV cells.

**Cells.** The Huh-7-derived H7-T7-IZ cells stably expressing the T7 RNA polymerase ([1]; kindly provided by Ralf Bartenschlager, University of Heidelberg, Germany) were maintained in a medium supplemented with 50 µg/ml of Zeocin. They were used for the transfection of the T7 promoter-driven pTM expression vectors. H7-T7-IZ cells were authenticated by Multiplex Cell Authentication (Multiplexion).

**Plasmids and transfection.** The pTM-ORF2-HEV-gt3 plasmid was kindly provided by J. Gouttenoire (University of Lausanne, Switzerland) [2]. The pTM-ORF2-HEV-gt1 plasmid was generated by cloning the Sar55 strain ORF2 sequence into the multiple cloning site of pTM plasmid. Plasmids were transfected into H7-T7-IZ cells using ViaFect™ Transfection Reagent (Promega) following the manufacturer's recommendations.

The plasmid pBlueScript SK(+)/HEV-p6 expressing a V5-tagged ORF1 (V1 insertion) has been described previously [3].

**Modeling of the N-terminus of ORF2**. We used an inhouse ColabFold implementation of AlphaFold2 [4] to generate models of the N-terminus of ORF2. Briefly, we used the ORF2 sequence or only the sequence encompassing up to residue 311 (comprising the shell (S) domain) or up to residue 446 (with the middle (M) domain added), with or without the 13 N-terminal residues (*i.e.* full-length or OR2i), to query the UniRef30 database (March or June 2021 release) with either HHBlits of MMseqs2 [5]. The resulting sequence alignments typically comprised 180-500 hits with 25-400 sequences aligned to any residue of the query. These alignments were used as inputs to AlphaFold2 and 5 models were generated for each. Resulting models were aligned to the S domain of PDB 3iyo. Molecules A, B and C, the three distinct positions in the T=3 icosahedral HEV capsid, were tried. This allowed displaying our models in the context of the capsid-like particle visualized by low resolution cryo-electron microscopy (cryo-EM, Electron Microscopy Data Bank entry 5173) [6]. The models and EM map were displayed and rendered with the PyMOL Molecular Graphics System (version 1.8 2015).

**Patient samples.** Patient samples were collected in France. This was a non-interventional study. Samples were obtained only *via* standard viral diagnostics following a physician's order (no supplemental or modified sampling). Data were analyzed anonymously. According to the French law (Loi Jardé), anonymous retrospective studies do not require institutional review board approval.

**RT-qPCR.** RNAs were extracted and next converted to cDNA by using a polydT primer and the AffinityScript Multiple Temperature cDNA Synthesis kit (Agilent Technologies). qPCR (TaqMan Gene Expression Assay, MGB-FAM-dye, ThermoFisher Scientific) were performed by using the QuantStudio3 Thermocycler (Applied Biosystems) and using primers (5'-GGTGGTTTCTGGGGTGAC-3' (F) and 5'-AGGGGTTGGTTGGATGAA-3' (R)) and a probe (5'-FAM-TGATTCTCAGCCCTTCGC-TAMRA-3') that target a conserved 70 bp region in the ORF2/3 overlap [7]. For cells harboring the subgenomic p6-replicon, qPCR were performed using primers (5'-AAGACATTCTGCGCTTTGTT-3' (F) and 5'-TGACTCCTCATAAGCATCGC-3' (R)) and a probe (5'-FAM-CCGTGGTTCCGTGCCATTGA-3') that target the ORF1 [8].

**Western blotting analyses.** Samples were separated by 10% SDS-PAGE and transferred onto nitrocellulose membranes (Hybond-ECL, Amersham). The targeted proteins were detected with specific antibodies (**Supplementary Table 1)** and corresponding peroxidase-conjugated secondary antibodies. The detection of proteins was done by chemiluminescence analysis (ECL, Amersham).

**Indirect immunofluorescence.** Cells were grown on coverslips in 24-well plates and fixed at 6 d.p.e. for electroporated cells and 12 d.p.i. for infected cells, with 3% of Paraformaldehyde (PFA). After 20 minutes (min), cells were washed twice with phosphate-buffered saline (PBS) and permeabilized for 5 min with cold methanol and then with 0.5% Triton X-100 (TX) for 30 min. Cells were incubated in PBS containing 10% goat serum for 30 min at room temperature (RT) and stained with the indicated primary antibodies for 30 min at RT followed by fluorochrome-conjugated secondary antibodies for 20 min at RT. The nuclei were stained with DAPI (4',6-diamidino-2-phenylindole). After 2 washes with PBS, coverslips were mounted with Mowiol 4-88 (Calbiochem) on glass slides and analyzed with a LSM 880 confocal laser-scanning microscope (Zeiss) using a Plan Apochromat 63xOil/1.4N.A. objective. The images were processed using Fiji software.

**Transferrin endocytosis.** PLC3/HEV and PLC3/HEV-ΔORF3 cells were incubated with 25 µg/ml of Alexa633-conjugated transferrin (TrF) at 37°C and fixed at indicated times. Cells were next permeabilized with methanol and TX, then stained with the P1H1 antibody. Cells were analyzed by confocal microscopy.

**Infectious titers**

Huh 7.5 cells were seeded in 96-well plates. The following day, cells were infected with serial dilutions of supernatants or intracellular viral particles from PLC3/HEV cells. Three days post-infection, cells were fixed and processed for indirect immunofluorescence. Cells labeled with anti-ORF2 antibody 1E6 were counted as infected cells. The number of infected cells was determined for each dilution and used to define the infectious titers in focus forming unit (FFU)/ml.

### Intracellular viral particles

Confluent T75 flasks of PLC3/HEV cells were trypsinized and cells were centrifuged for 10 min at 1500 rpm. Cells were washed thrice with PBS. Intracellular viral particles were extracted by resuspending cells in 1ml of sterile water at room temperature. Cells were vortexed vigorously for 20 min and then 110µl of sterile 10X PBS were added. Samples were clarified by centrifugation 2 min at 14000 rpm. The supernatants containing intracellular particles were collected and stored at -80°C until use.

### *In situ* labeling of viral RNA

PLC3 cells electroporated with HEV-p6 (PLC3/HEV) or HEV-p6 expressing a V5-tagged ORF1 (PLC3/HEV-ORF1V5) strains were fixed in 3% PFA for 20 min. Coverslips holding the fixed cells were attached to glass slides with a drop of nail polish and hydrophobic barriers were drawn around them with the ImmEdge Hydrophobic Barrier Pen (ACD Bio). Next, fixed cells were pre-treated according to the supplier instructions (RNAscope® $H_2O_2$ and Protease Reagents). First, the cells were treated with $H_2O_2$ for 10 min at RT and then washed twice with 1×PBS. Next, the protease III was diluted 1:15 and incubated for 15 min at RT; slides were washed twice. Then, the RNAscope assay was carried out following the user manual precisely (RNAscope Detection Kit Multiplex Fluorescent Reagent Kit v2 [9,10]. We used a probe (ref. 1030631-C2, Advanced Cell Diagnostics Bio-Techne) that targets positive strand of genomic viral RNAs [3]. The RNAs were labeled with fluorophore Opal 520 (Akoya Biosciences). Subsequently, immunofluorescent labeling using either P1H1, anti-ORF3 or anti-Rab11 antibodies was performed. Finally, coverslips were mounted and cells were analyzed by confocal microscopy.

**Supplementary Figure legends**

**Supplementary Figure 1: ORF2 and ORF3 antibody recognition in PLC3 mock cells.** Mock electroporated PLC3 cells were fixed, permeabilized with methanol and TX-0.5% and double-stained with indicated anti-ORF2 and anti-ORF3 antibodies. Red = ORF2; Green = ORF3; Blue = DAPI. Staining were analyzed by confocal microscopy. Scale bar, 20μm.

**Supplementary Figure 2: Antibody recognition of gt1 and gt3 ORF2 proteins.** H7-T7-IZ cells transfected with pTM plasmid expressing p6 strain ORF2 (ORF2-gt3) or Sar55 strain ORF2 (ORF2-gt1) were fixed at 16h post-transfection, permeabilized with Methanol and TX-0.5% and stained with indicated anti-ORF2 antibodies. Staining were analyzed by confocal microscopy. Red = ORF2; Blue = DAPI. Scale bar, 20μm.

**Supplementary Figure 3: Immunogold labeling of PLC3 and Huh-7.5 mock cells.** Cryosections of PLC3 and Huh-7.5 mock cells were immunogold-labeled with the indicated antibodies and analyzed by EM.

**Supplementary Figure 4: Identification of HEV-induced vesicular and tubular structures in HEV-producing Huh-7.5 cells.** Huh-7.5 cells electroporated with HEV RNA (**A-C**) and Huh-7.5 cells infected with HEV particles (**D-F**) were fixed at 6 days p.e and 12 days post-infection, respectively. (**A, D**) Cells were next processed for immunostaining with 1E6, P1H1, anti-ORF3 and anti-Rab11 antibodies, as indicated. Red = ORF2; Green = ORF3 or Rab11; Blue = DAPI. Staining were analyzed by confocal microscopy. Scale bar, 20μm. Manders' Overlap Coefficients (MOC) of ORF2i

(P1H1) staining in ORF3 or Rab11 staining ($n \geq 30$ cells) were calculated. (**B-C, E-F**) Cryosections of HEV-producing Huh-7.5 cells were processed for immunogold labeling with 1E6 or P1H1 antibodies (visualized by 6 nm gold particles), as indicated. Cryosections were next analyzed by EM. Vesicular (**B, E**) and Tubular (**C, F**) structures containing ORF2 proteins are indicated by black arrows. N, nucleus.

**Supplementary Figure 5: Colocalization analysis of the ORF2i protein with different cell markers in PLC3/HEV cells.** PLC3/HEV cells were fixed, permeabilized with cold methanol and TX-0.5% and double-stained with P1H1 and anti-cell marker antibodies, as indicated. Staining were analyzed by confocal microscopy. Scale bar, 20μm.

**Supplementary Figure 6: Colocalization analysis of the ORF2i protein with different cell markers in PLC3 mock cells.** PLC3 mock cells were fixed, permeabilized with methanol and TX-0.5% and double-stained with P1H1 and anti-cell marker antibodies, as indicated. Staining were analyzed by confocal microscopy. Scale bar, 20μm.

**Supplementary Figure 7: Kinetics of colocalization of the ORF2i protein with transferrin.** PLC3/HEV (**A**) and PLC3/HEV-ΔORF3 (**B**) cells were incubated with fluorochrome-conjugated transferrin (TrF) at 37°C and fixed at indicated times. Cells were next permeabilized and stained with the P1H1 antibody. Red = ORF2; Green = transferrin; Blue = DAPI. Staining were analyzed by confocal microscopy. Scale bar, 20μm. (**B**) Manders' Overlap Coefficients (MOC) of ORF2 staining in TrF staining ($n \geq$ 50 cells).

**Supplementary Figure 8: Double-immunogold labeling of PLC3 mock cells.** Cryosections of PLC3 mock cells were processed for double immunogold labeling with anti-ORF2 or anti-ORF3 (visualized by 10 nm gold particles) and anti-Rab11, anti-CD71 or anti-CD81 (visualized by 6 nm gold particles) antibodies, as indicated. Cryosections were next analyzed by EM.

**Supplementary Figure 9: Detection of ORF1 protein and genomic HEV RNA in HEV-producing PLC3 cells. (A)** PLC3 cells were electroporated with the p6 strain expressing either wildtype (PLC3/HEV) or the V5-tagged ORF1 (PLC3/HEV-ORF1V5) proteins or mock electroporated (PLC3 mock). At 3 d.p.e., cells were processed for immunofluorescence using anti-V5 (V5, red), and P1H1 (ORF2, green) or anti-ORF3 (ORF3, green) antibodies prior to analysis by confocal microscopy. (**B**) PLC3/HEV and PLC3 mock cells were grown on coverslips, fixed at 3 d.p.e. and processed for *in situ* RNAscope hybridization. Cells were stained with a probe targeting HEV genomic RNA (RNA, red) and anti-ORF2 (P1H1), anti-ORF3 or anti-Rab11 antibodies (green). Nuclei are in blue. Scale bar, 20$\mu$m.

**Supplementary Table 1: Primary antibodies used in Western-blot (WB), immunofluorescence (IF) and immunogold (IG) experiments.**

| Name | Target / Epitope | Host | Isotype | Source | Reference | Ab Registry | WB | IF | IG |
|---|---|---|---|---|---|---|---|---|---|
| P1H1 | ORF2i<br>GQPSGRRRGRRSGG | Mouse | IgG3 | This study | n/a | n/a | 1/500 | 1/500 | 1/100 |
| P2H1 | ORF2i<br>AGYPYNYNTTASDQ | Mouse | IgG2a | This study | n/a | n/a | 1/500 | 1/500 | 1/100 |
| P2H2 | ORF2i<br>AGYPYNYNTTASDQ | Mouse | IgG1 | This study | n/a | n/a | 1/500 | 1/500 | 1/100 |
| P3H2 | ORF2i/g/c<br>SRVVIQDYDNQHEQDR | Mouse | IgG3 | This study | n/a | n/a | 1/500 | 1/500 | 1/100 |
| 1E6 | ORF2i/g/c<br>GDSRVVIQDYDNQHEQ<br>DRPTPSPA | Mouse | IgG2b | Millipore | MAB8002 | AB_827236 | 1/2000 | 1/800 | 1/100 |
| ORF3 | ORF3<br>ANPPDHSAPLGVTRPSA<br>PPLPHVVDLPQLGPRR | Rabbit | pAb | S. Emerson | [11] | n/a | n/a | 1/1000 | 1/100 |
| Tub | $\beta$-tubulin<br>C-terminal region | Mouse | IgG1 | Sigma | T5201 | AB_609915 | 1/1000 | 1/100 | n/a |
| MTOC | $\gamma$-tubulin<br>N-terminal region | Mouse | IgG1 | Sigma | T5326 | AB_532292 | n/a | 1/500 | n/a |
| Calnexin | Human Calnexin | Rabbit | | Abcam | ab22595 | AB_2069006 | n/a | 1/1000 | n/a |
| ERGIC53 | ERGIC 53kDa protein | Mouse | IgG1 | Enzo Life Sciences Inc | ALX-804-602-C100 | AB_2051363 | n/a | 1/100 | n/a |
| EEA1 | Early Endosome Antigen 1 | Mouse | IgG1 | Transduction Laboratories | 610457 | AB_397830 | n/a | 1/500 | n/a |
| Rab5 | Rab5 | Rabbit | (mAb) | Cell signaling | #3547 | AB_2300649 | 1/1000 | 1/200 | n/a |
| Rab9a | Rab9a | Rabbit | (mAb) | Cell signaling | #5118 | AB_10621426 | n/a | 1/50 | n/a |
| Rab11 | Rab11 | Rabbit | (mAb | Cell signaling | #5589 | AB_10693925 | n/a | 1/50 | n/a |
| Rab11a | Rab11a | Rabbit | (pAb) | ThermoFisher | 71-5300 | AB_2533987 | n/a | n/a | 1/10 |
| Rab11a | Rab11a | Rabbit | (mAb) | Abcam | ab128913 | AB_11140633 | 1/20000 | n/a | n/a |
| Rab11b | Rab11b | Rabbit | (mAb) | Abcam | ab175925 | n/a | 1/10000 | n/a | n/a |
| 5A6 | Tetraspanin CD81<br>Large extracellular loop | Mouse | IgG1 | S. Levy | [12] | AB_627192 | n/a | 1/300 | 1/30 |
| CD63 | Tetraspanin CD63<br>Large extracellular loop | Mouse | IgG1 | BD Pharmingen | 556019 | AB_396297 | n/a | 1/500 | n/a |
| CD71 | Transferrin receptor | Mouse | IgG1 | Santa Cruz Biotechnology | sc-65882 | AB_1120670 | n/a | 1/100 | n/a |
| CD71 | Transferrin receptor | Rabbit | pAb | Abcam | ab84036 | AB_10673794 | n/a | 1/1000 | 1/100 |
| EHD1 | Eps15 homology domain protein 1 | Rabbit | (mAb) | Abcam | ab109747 | AB_10864800 | n/a | 1/1000 | n/a |
| MICAL-L1 | Molecule Interacting with CasL-like1 | Rabbit | IgG | Abcam | ab220648 | n/a | n/a | 1/100 | n/a |
| PACSIN2 | Protein Kinase C and Casein Kinase Substrate in Neurons 2 | Rabbit | IgG | MyBiosource | MBS7114698 | n/a | n/a | 1/1000 | n/a |
| PMP70 | 70-kDa Peroxisomal Membrane Protein | Mouse | IgG1 | Sigma | SAB4200181 | AB_10639362 | n/a | 1/1000 | n/a |
| Catalase | Catalase | Rabbit | IgG | Cell signaling | #12980 | AB_2798079 | n/a | 1/800 | n/a |
| TOM-20 | Translocase of the outer mitochondrial Membrane | Mouse | IgG1 | BD Biosciences | 612278 | AB_399595 | n/a | 1/100 | n/a |
| LAMP1 | Lysosomal associated membrane protein 1 | Rabbit | (mAb) | Cell signaling | #9091 | AB_2687579 | n/a | 1/200 | n/a |
| V5 | GKPIPNPLLGLDST | Mouse | IgG2a | Abcam | ab27671 | AB_471093 | n/a | 1/500 | n/a |

**References**


[1] Romero-Brey I, Merz A, Chiramel A, Lee J-Y, Chlanda P, Haselman U, et al. Three-dimensional architecture and biogenesis of membrane structures associated with hepatitis C virus replication. PLoS Pathogens 2012;8:e1003056. https://doi.org/10.1371/journal.ppat.1003056.

[2] Lenggenhager D, Gouttenoire J, Malehmir M, Bawohl M, Honcharova-Biletska H, Kreutzer S, et al. Visualization of hepatitis E virus RNA and proteins in the human liver. Journal of Hepatology 2017;67:471–9. https://doi.org/10.1016/j.jhep.2017.04.002.

[3] Metzger K, Bentaleb C, Hervouet K, Alexandre V, Montpellier C, Saliou J-M, et al. Processing and Subcellular Localization of the Hepatitis E Virus Replicase: Identification of Candidate Viral Factories. Front Microbiol 2022;13:828636. https://doi.org/10.3389/fmicb.2022.828636.

[4] Jumper J, Evans R, Pritzel A, Green T, Figurnov M, Ronneberger O, et al. Highly accurate protein structure prediction with AlphaFold. Nature 2021;596:583–9. https://doi.org/10.1038/s41586-021-03819-2.

[5] Gabler F, Nam S, Till S, Mirdita M, Steinegger M, Söding J, et al. Protein Sequence Analysis Using the MPI Bioinformatics Toolkit. Curr Protoc Bioinform 2020;72:e108. https://doi.org/10.1002/cpbi.108.

[6] Xing L, Li TC, Mayazaki N, Simon MN, Wall JS, Moore M, et al. Structure of hepatitis E virion-sized particle reveals an RNA-dependent viral assembly pathway. Journal of Biological Chemistry 2010;285:33175–83. https://doi.org/10.1074/jbc.m110.106336.

[7] Jothikumar N, Cromeans TL, Robertson BH, Meng XJ, Hill VR. A broadly reactive one-step real-time RT-PCR assay for rapid and sensitive detection of hepatitis E virus. Journal of Virological Methods 2006;131:65–71. https://doi.org/10.1016/j.jviromet.2005.07.004.

[8] Yin X, Ying D, Lhomme S, Tang Z, Walker CM, Xia N, et al. Origin, antigenicity, and function of a secreted form of ORF2 in hepatitis E virus infection. Proceedings of the National Academy of Sciences of the United States of America 2018;3:201721345–6. https://doi.org/10.1073/pnas.1721345115.

[9] Wang F, Flanagan J, Su N, Wang L-C, Bui S, Nielson A, et al. RNAscope A Novel in Situ RNA Analysis Platform for Formalin-Fixed, Paraffin-Embedded Tissues. J Mol Diagnostics 2012;14:22–9. https://doi.org/10.1016/j.jmoldx.2011.08.002.

[10] Liu D, Tedbury PR, Lan S, Huber AD, Puray-Chavez MN, Ji J, et al. Visualization of Positive and Negative Sense Viral RNA for Probing the Mechanism of Direct-Acting Antivirals against Hepatitis C Virus. Viruses 2019;11:1039. https://doi.org/10.3390/v11111039.

[11] Graff J, Nguyen H, Yu C, Elkins WR, Claire MS, Purcell RH, et al. The Open Reading Frame 3 Gene of Hepatitis E Virus Contains a cis-Reactive Element and Encodes a Protein Required for Infection of Macaques. Journal of Virology 2005;79:6680–9. https://doi.org/10.1128/jvi.79.11.6680-6689.2005.

[12] Oren R, Takahashi S, Doss C, Levy R, Levy S. TAPA-1, the target of an antiproliferative antibody, defines a new family of transmembrane proteins. Molecular and Cellular Biology 1990;10:4007. https://doi.org/10.1128/mcb.10.8.4007.

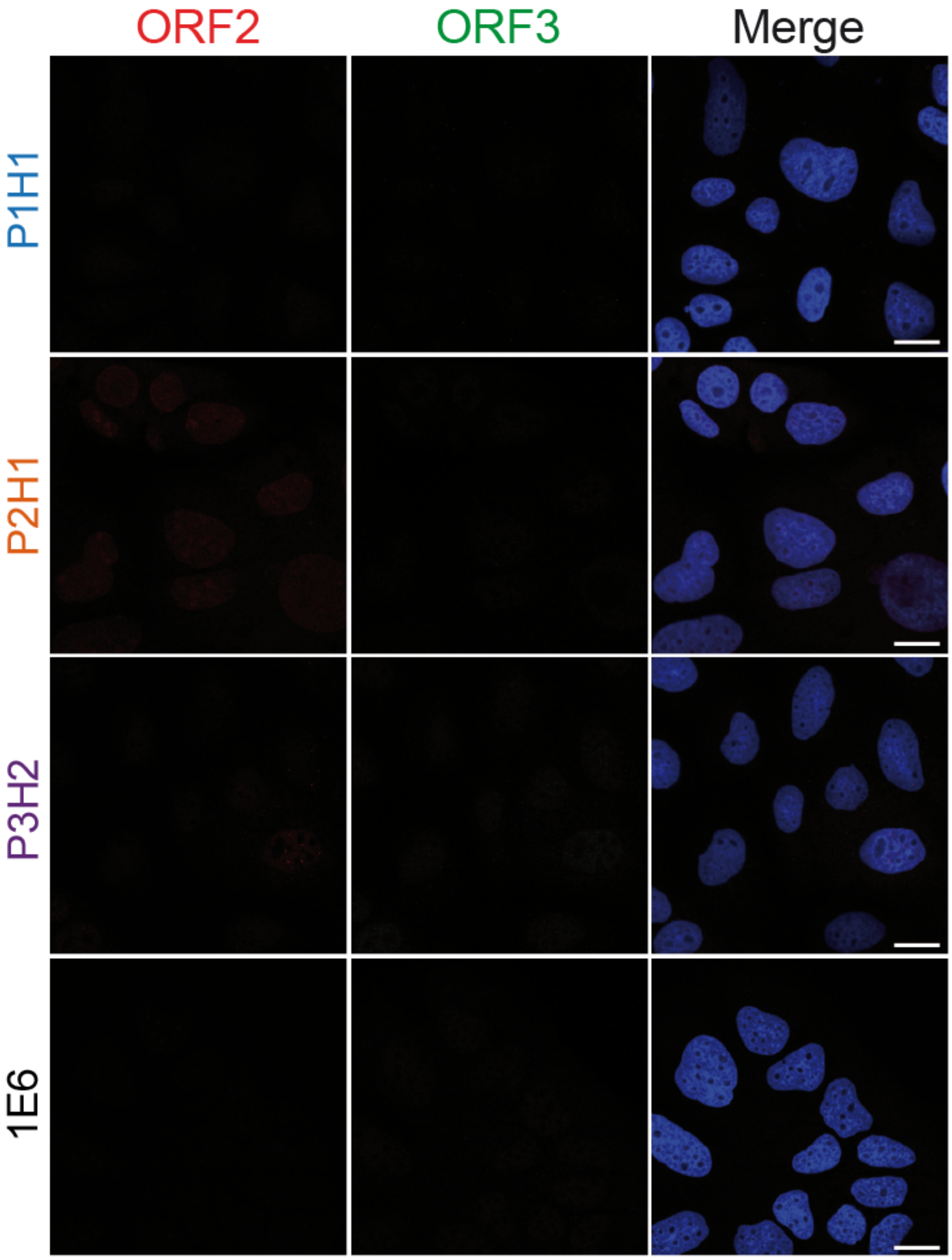


**Fig S1**

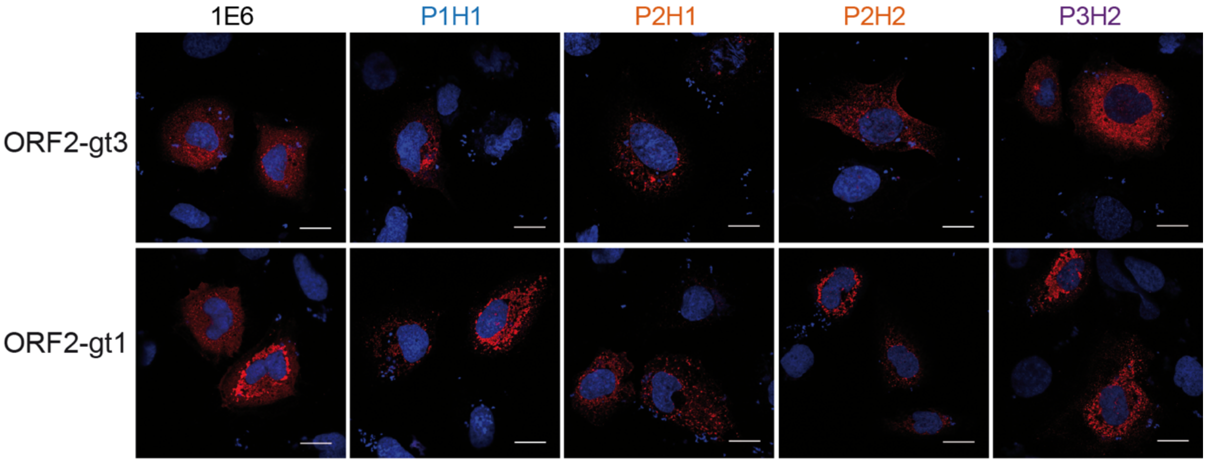


**Fig S2**

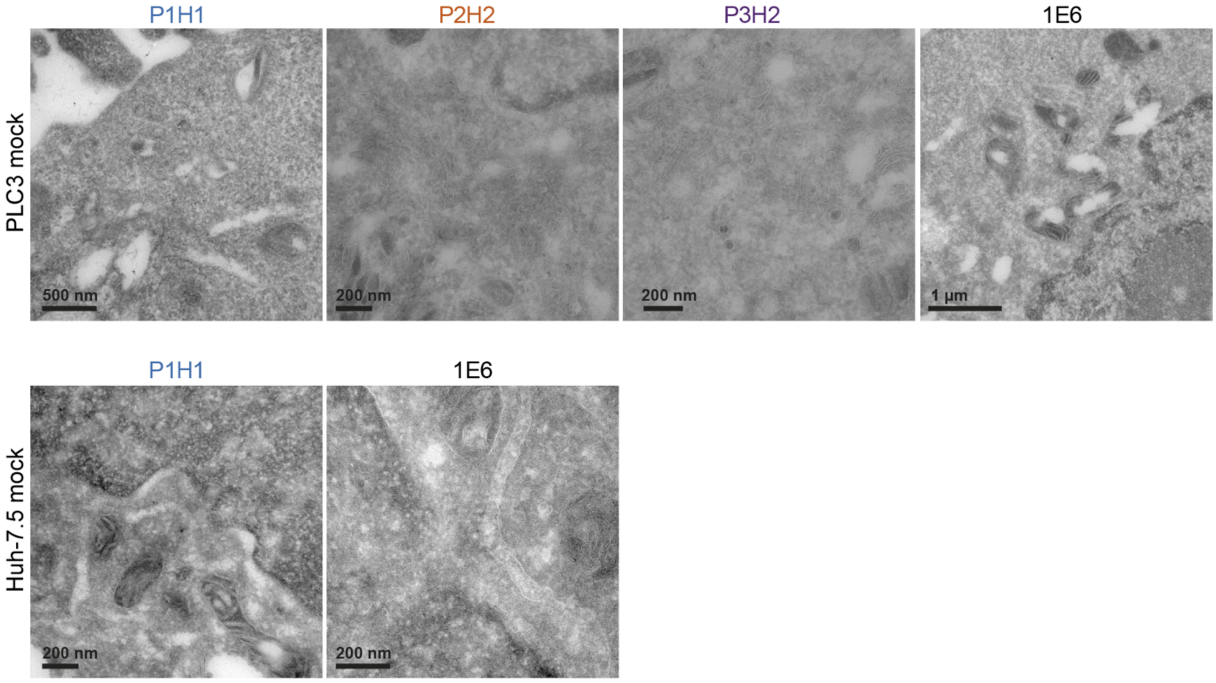

P1H1
P2H2
P3H2
1E6
PLC3 mock
500 nm
200 nm
200 nm
1 µm
P1H1
1E6
Huh-7.5 mock
200 nm
200 nm


**Fig S3**

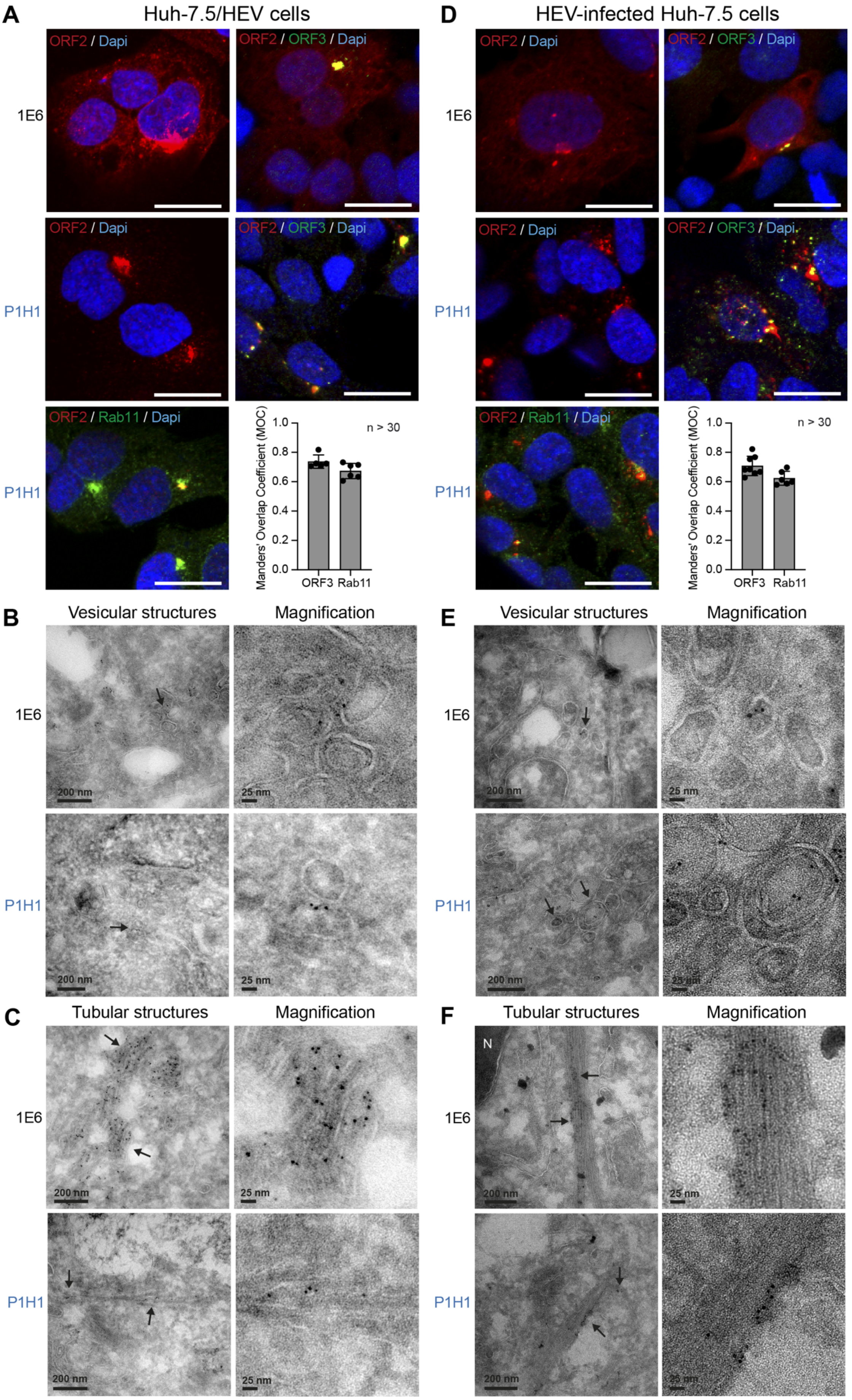
A
Huh-7.5/HEV cells
ORF2 / Dapi
ORF2 / ORF3 / Dapi
1E6
P1H1
ORF2 / Rab11 / Dapi
Manders' Overlap Coefficient (MOC)
n > 30
ORF3
Rab11
D
HEV-infected Huh-7.5 cells
B
Vesicular structures
Magnification
200 nm
25 nm
E
C
Tubular structures
F
N

**Fig S4**

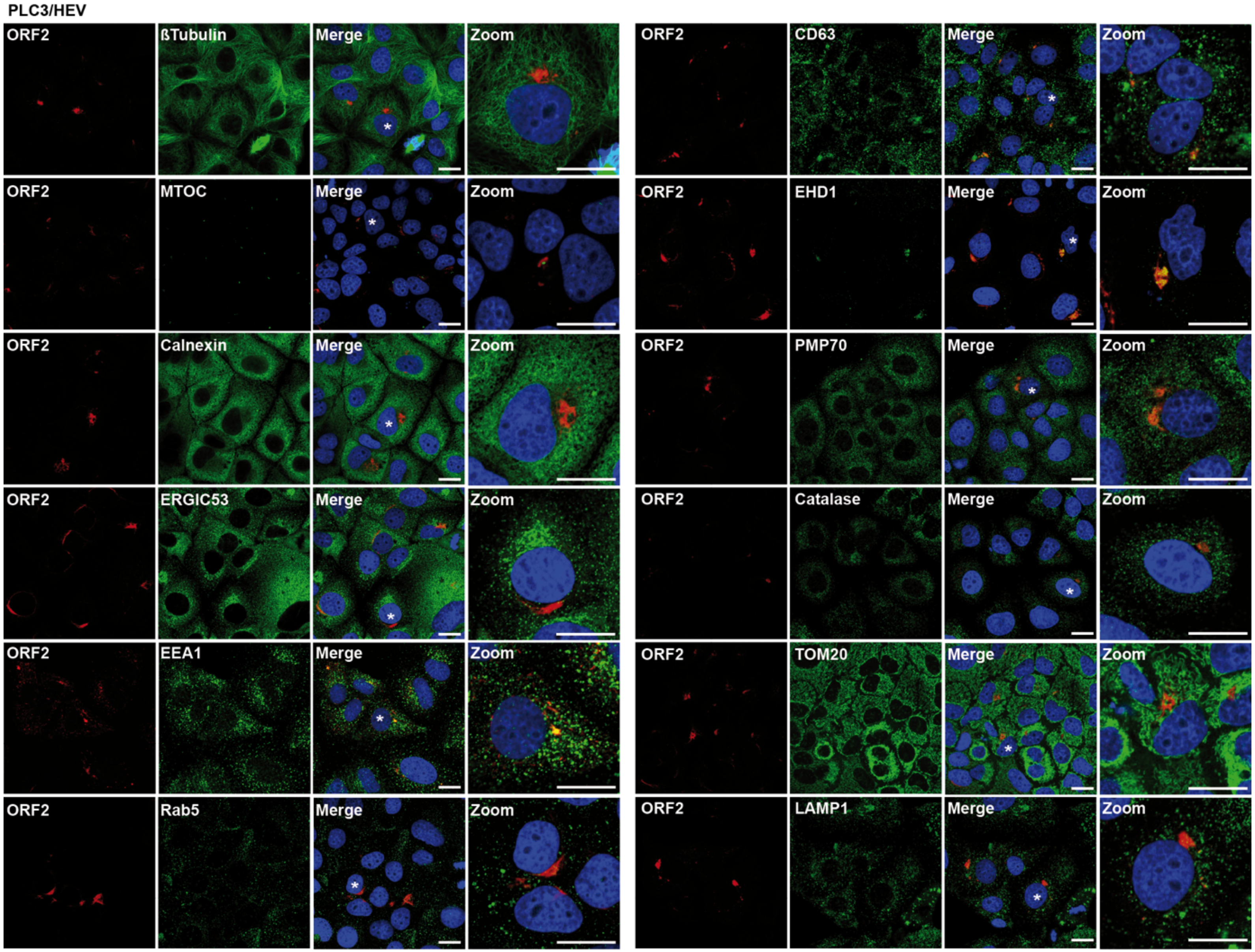
PLC3/HEV
ORF2
ßTubulin
Merge
Zoom
ORF2
CD63
Merge
Zoom
ORF2
MTOC
Merge
Zoom
ORF2
EHD1
Merge
Zoom
ORF2
Calnexin
Merge
Zoom
ORF2
PMP70
Merge
Zoom
ORF2
ERGIC53
Merge
Zoom
ORF2
Catalase
Merge
Zoom
ORF2
EEA1
Merge
Zoom
ORF2
TOM20
Merge
Zoom
ORF2
Rab5
Merge
Zoom
ORF2
LAMP1
Merge
Zoom

Fig S5

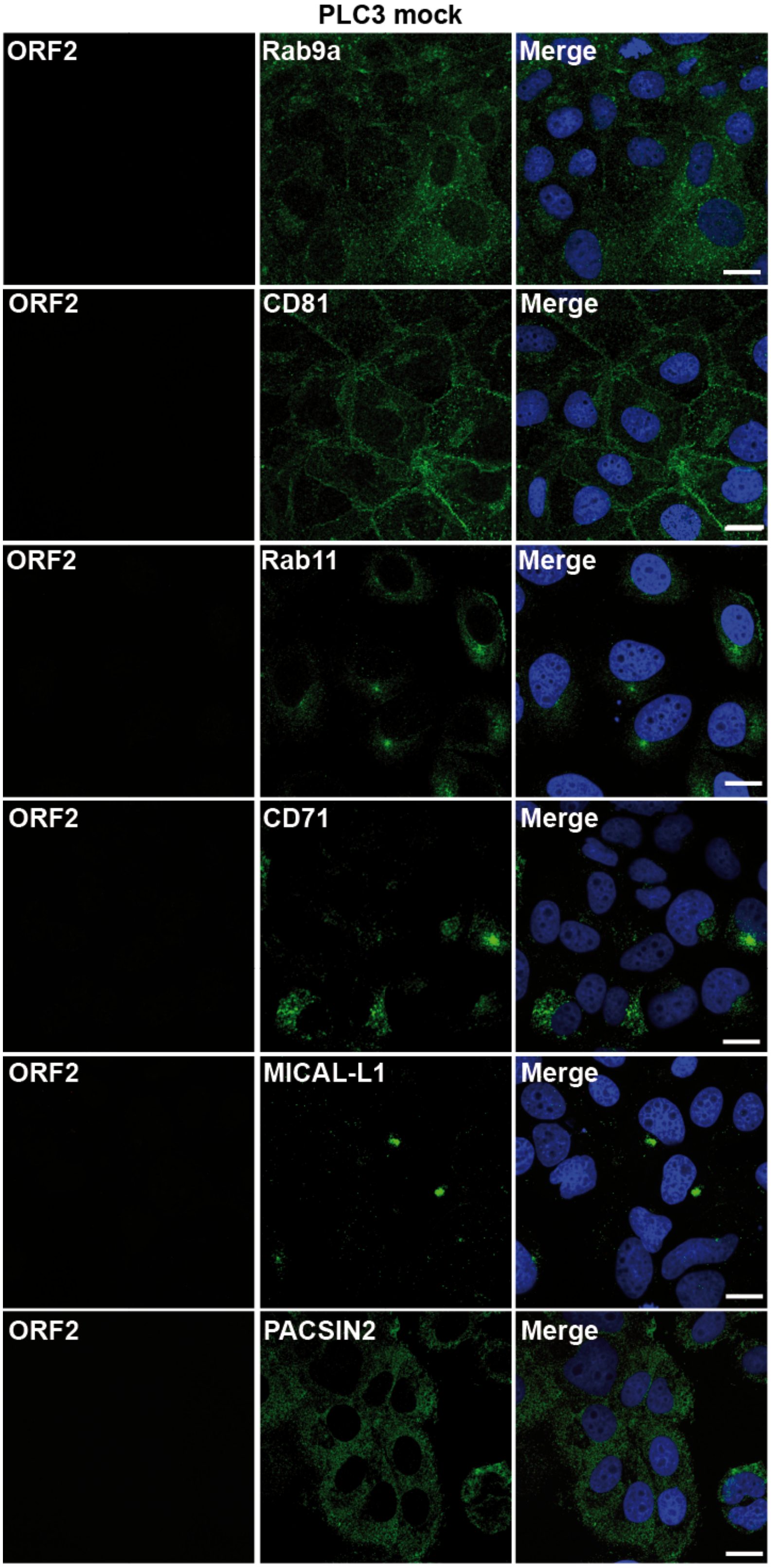
PLC3 mock
ORF2
Rab9a
Merge
ORF2
CD81
Merge
ORF2
Rab11
Merge
ORF2
CD71
Merge
ORF2
MICAL-L1
Merge
ORF2
PACSIN2
Merge

**Fig S6**

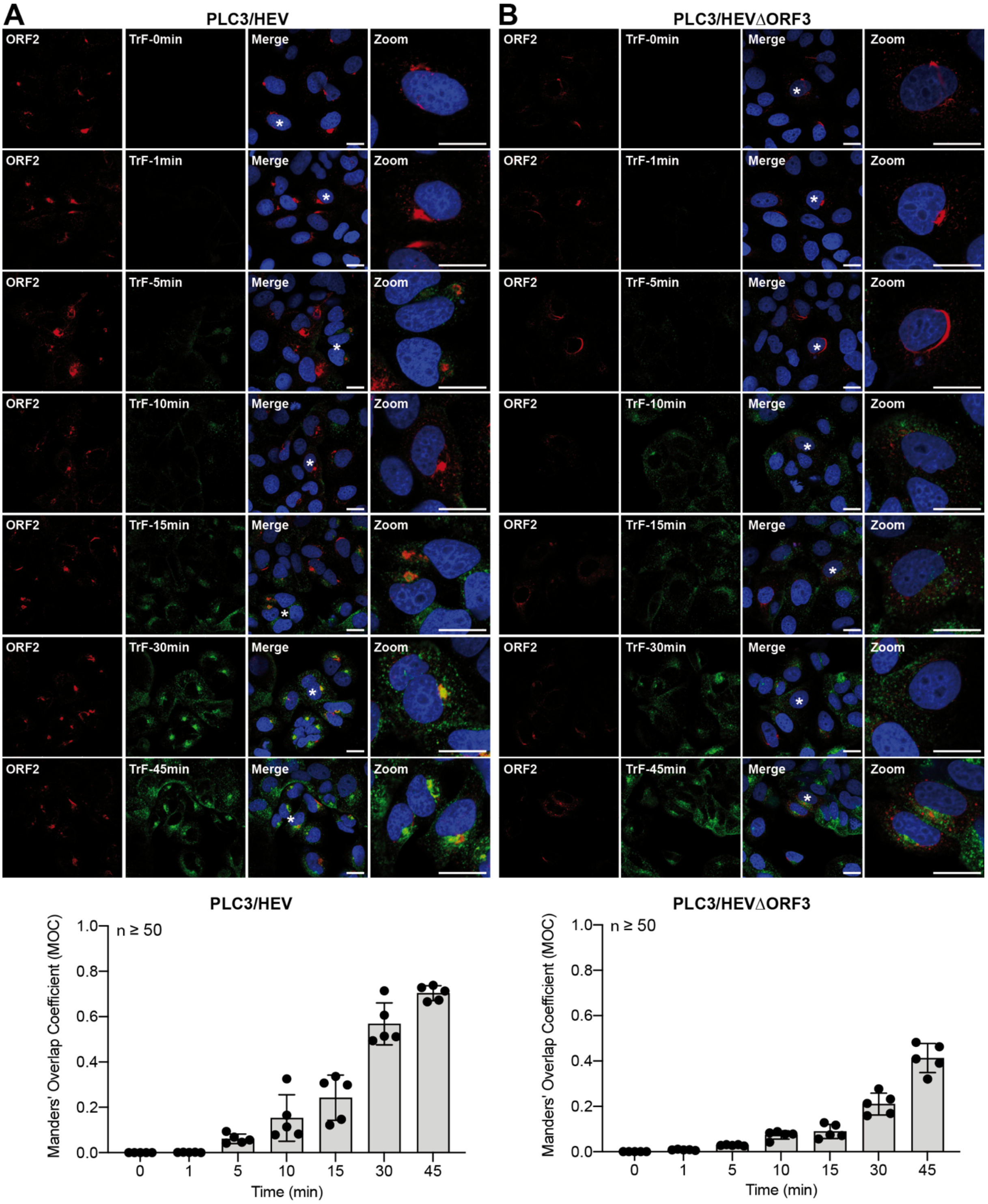
A
PLC3/HEV
B
PLC3/HEVΔORF3
ORF2
TrF-0min
TrF-1min
TrF-5min
TrF-10min
TrF-15min
TrF-30min
TrF-45min
Merge
Zoom
PLC3/HEV
PLC3/HEVΔORF3
n ≥ 50
Manders' Overlap Coefficient (MOC)
1.0
0.8
0.6
0.4
0.2
0.0
0
1
5
10
15
30
45
Time (min)

**Fig S7**

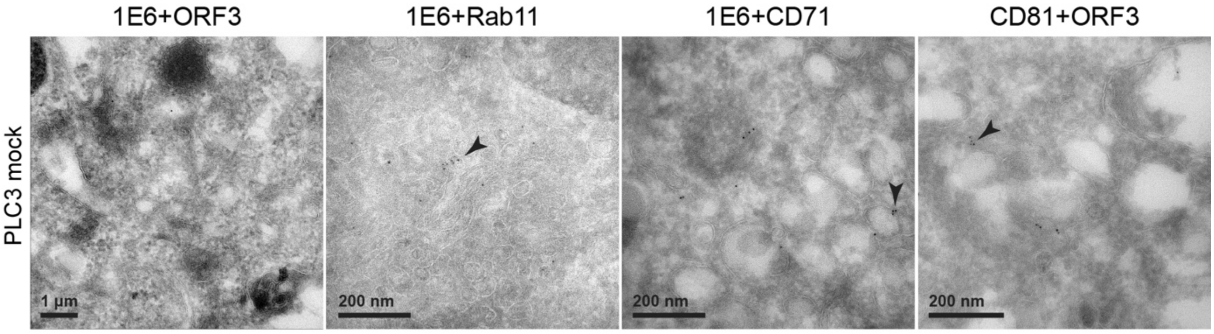

1E6+ORF3
1E6+Rab11
1E6+CD71
CD81+ORF3
PLC3 mock
1 µm
200 nm
200 nm
200 nm


**Fig S8**

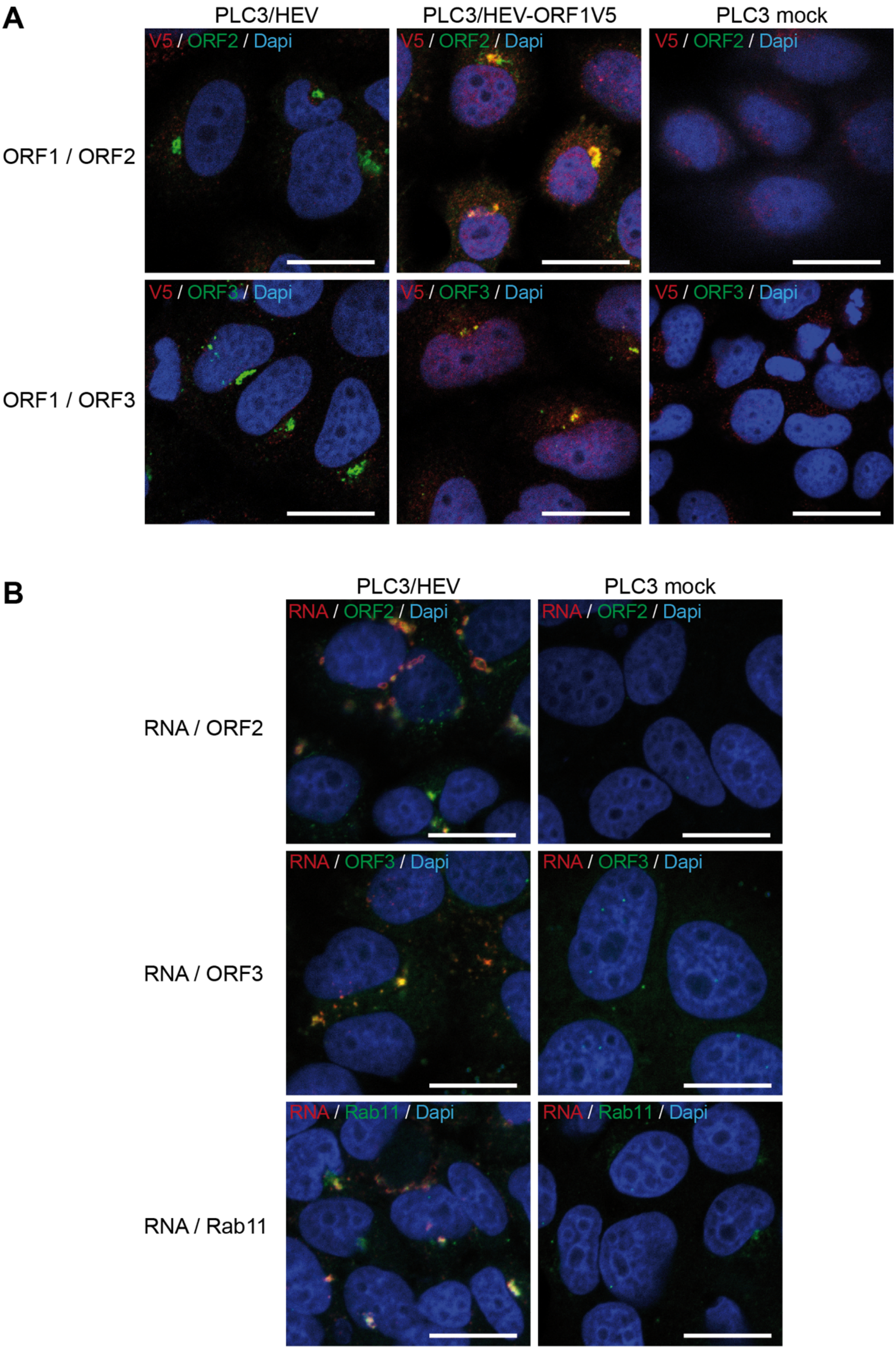
A
PLC3/HEV
PLC3/HEV-ORF1V5
PLC3 mock
V5 / ORF2 / Dapi
ORF1 / ORF2
V5 / ORF3 / Dapi
ORF1 / ORF3
B
PLC3/HEV
PLC3 mock
RNA / ORF2 / Dapi
RNA / ORF2
RNA / ORF3 / Dapi
RNA / ORF3
RNA / Rab11 / Dapi
RNA / Rab11

**Fig S9**
